# Properties of Glial Cell at the Neuromuscular Junction are Incompatible with synaptic repair in the SOD1^G37R^ ALS mouse model

**DOI:** 10.1101/2020.06.05.136226

**Authors:** Martineau Éric, Danielle Arbour, Joanne Vallée, Robitaille Richard

**Affiliations:** Département de neurosciences, Université de Montréal, PO box 6128, Station centre-ville, Montréal, Québec, Canada, H3C 3J7; Groupe de recherche sur le système nerveux central, Université de Montréal

**Author notes:** To whom correspondence should be addressed, Université de Montréal, PO box 6128, Station centre-ville, Montréal, Québec, Canada, H3C 3J7.

## Abstract

Amyotrophic lateral sclerosis (ALS) is a fatal neurodegenerative disease affecting motoneurons in a motor-unit (MU) dependent manner. Glial dysfunction contributes to numerous aspects of the disease. At the neuromuscular junction (NMJ), early alterations in perisynaptic Schwann cell (PSC), glial cells at this synapse, may impact their ability to regulate NMJ stability and repair. Indeed, muscarinic receptors (mAChR) regulate the repair phenotype of PSCs and are overactivated at disease-resistant NMJs (*Soleus* muscle) in SOD1^G37R^ mice. However, it remains unknown whether this is the case at disease-vulnerable NMJs and whether it translates into an impairment of PSC-dependent repair mechanisms. We used *Soleus* and *Sternomastoid* muscles from SOD1^G37R^ mice and performed Ca^2+^-imaging to monitor PSC activity and used immunohistochemistry to analyze their repair and phagocytic properties. We show that PSC mAChR-dependent activity was transiently increased at disease-vulnerable NMJs (*Sternomastoid* muscle). Furthermore, PSCs from both muscles extended disorganized processes from denervated NMJs and failed to initiate or guide nerve terminal sprouts at disease-vulnerable NMJs, a phenomenon essential for compensatory reinnervation. This was accompanied by a failure of numerous PSCs to upregulate Galectin-3 (MAC-2), a marker of glial axonal debris phagocytosis, upon NMJ denervation in SOD1 mice. Finally, differences in these PSC-dependent NMJ repair mechanisms were MU-type dependent, thus reflecting MU vulnerability in ALS. Together, these results reveal that neuron-glia communication is ubiquitously altered at the NMJ in ALS. This appears to prevent PSCs from adopting a repair phenotype, resulting in a maladapted response to denervation at the NMJ in ALS.

**SIGNIFICANCE STATEMENT:** Understanding how the complex interplay between neurons and glial cells ultimately lead to the degeneration of motor neurons and loss of motor function is a fundamental question to comprehend amyotrophic lateral sclerosis. An early and persistent alteration of glial cell activity takes place at the neuromuscular junction (NMJ), the output of motor neurons, but its impact on NMJ repair remains unknown. Here, we reveal that glial cells at disease-vulnerable NMJs often fail to guide compensatory nerve terminal sprouts and to adopt a phagocytic phenotype on denervated NMJs in SOD1^G37R^ mice. These results show that glial cells at the NMJ elaborate an inappropriate response to NMJ degeneration in a manner that reflects motor-unit vulnerability and potentially impairs compensatory reinnervation.

## INTRODUCTION

Amyotrophic lateral sclerosis (ALS) is a neurodegenerative disease characterized by the progressive loss of motoneurons (MNs), with fast-fatigable (FF) motor-units (MUs) being more vulnerable than fast-fatigue-resistant (FR) and slow (S) MUs (Frey et al., 2000; Pun et al., 2006; Hegedus et al., 2007; Hegedus et al., 2008). Over the last decade, evidence of disease onset and progression being modulated by numerous non-cell autonomous mechanisms has highlighted the importance of glial cells in this disease (Boillee et al., 2006a; Ilieva et al., 2009; Kang et al., 2013; Ditsworth et al., 2017). Evidence from ALS patients and *SOD1* mouse models also shows early loss of neuromuscular junctions (NMJ) prior to disease onset (Fischer et al., 2004; Pun et al., 2006; Hegedus et al., 2007; Hegedus et al., 2008; Martineau et al., 2018), supporting the notion that ALS has a long silent pre-symptomatic phase (Eisen et al., 2014). Interestingly, an early and persistent functional alteration of perisynaptic Schwann cells (PSCs), glial cells at the NMJ, was reported in the *SOD1*^*G37R*^ mouse model (Arbour et al., 2015).

Early PSC dysfunction in ALS could be of particular importance owing to their ability to detect and modulate synaptic communication, regulate NMJ stability and oversee NMJ repair. Importantly, these three functions are interdependent as PSC synaptic decoding regulates their ability to stabilize or repair NMJs, notably through muscarinic receptors (mAChRs). Indeed, activation of PSC mAChRs and purinergic receptors triggers a transient increase in cytoplasmic calcium (Ca^2+^ response), through which they modulate neurotransmitter release (Robitaille, 1998; Rochon et al., 2001; Rousse et al., 2010; Todd et al., 2010). Blockade of PSC mAChRs destabilizes NMJs (Wright et al., 2009), demonstrating that proper PSC synaptic decoding is necessary for NMJ maintenance.

Importantly, PSCs contribute to NMJ re-innervation following denervation of NMJs by adopting a “pro-regenerative” (repair) phenotype. For instance, PSCs extend long processes known to guide re-innervation through nerve terminal sprouting (Reynolds and Woolf, 1992; Son and Thompson, 1995b, a; Son et al., 1996; O’Malley et al., 1999). Similarly to axonal Schwann cells (SCs) (Reichert et al., 1994; Painter et al., 2014; Brosius Lutz et al., 2017), PSCs also actively phagocytose presynaptic debris following denervation (Duregotti et al., 2015) which is essential for efficient axonal regrowth and NMJ re-innervation (Kang and Lichtman, 2013). Finally, complete PSC endplate coverage is essential for re-innervation since parts of the endplate vacated by PSCs are forsaken during re-innervation (Kang et al., 2014). Importantly, mAChRs activation represses PSC’s repair phenotype and can modulate NMJ repair by regulating PSC gene expression (Georgiou et al., 1994; Georgiou et al., 1999).

Interestingly, Arbour et al. (2015) reported an enhanced mAChR contribution to PSC Ca^2+^ responses at NMJs innervated by disease-resistant or moderately vulnerable MUs (S and FR MU respectively, *Soleus* (SOL) muscle) in *SOD1*^*G37R*^ mice. As previously proposed (Ko and Robitaille, 2015; Arbour et al., 2017), we postulated that inappropriate mAChR signaling could prevent PSCs from adopting a repair phenotype, thus hindering NMJ reinnervation in ALS. However, this repair phenotype remains unevaluated in symptomatic animals. Furthermore, recent evidence revealed opposite changes in the synaptic properties of NMJs from different MU types in *SOD1*^*G37R*^ mice (Tremblay et al., 2017). This raises the possibility that PSC properties, notably mAChR overactivation, may be different at NMJs innervated by vulnerable FF MUs.

In the present study, we assessed if PSCs mAChR overactivation was also present at NMJs innervated by vulnerable FF MUs in the *Sternomastoid* (STM) muscle at different stages of the disease. Next, we evaluated if PSCs adopted a repair phenotype, either in presymptomatic SOD1^G37R^ animals following experimentally-induced denervation, or in symptomatic SOD1^G37R^ mice where the progressive denervation of NMJs is ongoing. Notably, we evaluated the presence of extended processes, nerve terminal sprouts and phagocytic markers at vulnerable (STM or EDL) or partially resistant (SOL) NMJs. Albeit through a different mechanism, PSCs in the STM also displayed an early, but transient, increase in mAChR-dependent decoding. Consistent with defects in PSC-dependent NMJ repair mechanisms in all these muscles, PSCs displayed abnormal process extension as well as paradoxical expression of Galectin-3 (MAC-2), a marker of glial phagocytosis. Altogether, these results suggest that changes in PSCs are maladapted in ALS.

## MATERIALS AND METHODS

### Animals

Male transgenic mice heterozygote for the human *Sod1* gene carrying the *G37R* mutation (*SOD1*^*G37R*^, line 29; [B6.Cg-Tg(SOD1*G37R)29Dpr/J]; stock number 008229), or, in some experiments, the *G93A* mutation (*SOD1*^*G93A*^; [B6SJL-Tg(SOD1*G93A)1Gur/J]; stock number 002726), were obtained from The Jackson laboratories (Bar Harbor, ME). *SOD1*^*G37R*^ mice were maintained on a C57BL/6 background and were genotyped by PCR for the *hSOD1* gene performed on a tail sample taken at the time of the weaning.

For some experiments, transgenic mice expressing the “yellow fluorescent protein” (YFP) in all motor neurons (Feng et al., 2000) were used (homozygous *Thy1-YFP*, line 16; [B6.Cg-Tg(Thy1-YFP)16Jrs/J]; stock number 003709). These mice were also obtained from The Jackson laboratories (Bar Harbor, ME) and were maintained on a C57BL/6 background.

Pre-symptomatic *SOD1*^*G37R*^ mice were used at P119-131 (P120) and P179-194 (P180) (Figure 1A). Phenotypically matched late symptomatic *SOD1*^*G37R*^ mice were used between P505 and P565. Age-matched *wild-type (WT*) brothers were used as controls. Animals were sacrificed by exposition to a lethal dose of isoflurane (CDMV, Saint-Hyacinthe, Qc, Canada, J2S 7C2). All experiments were performed in accordance with the guidelines of the Canadian Council of Animal Care and the Comité de déontologie animale of Université de Montréal.

**Figure 1:**
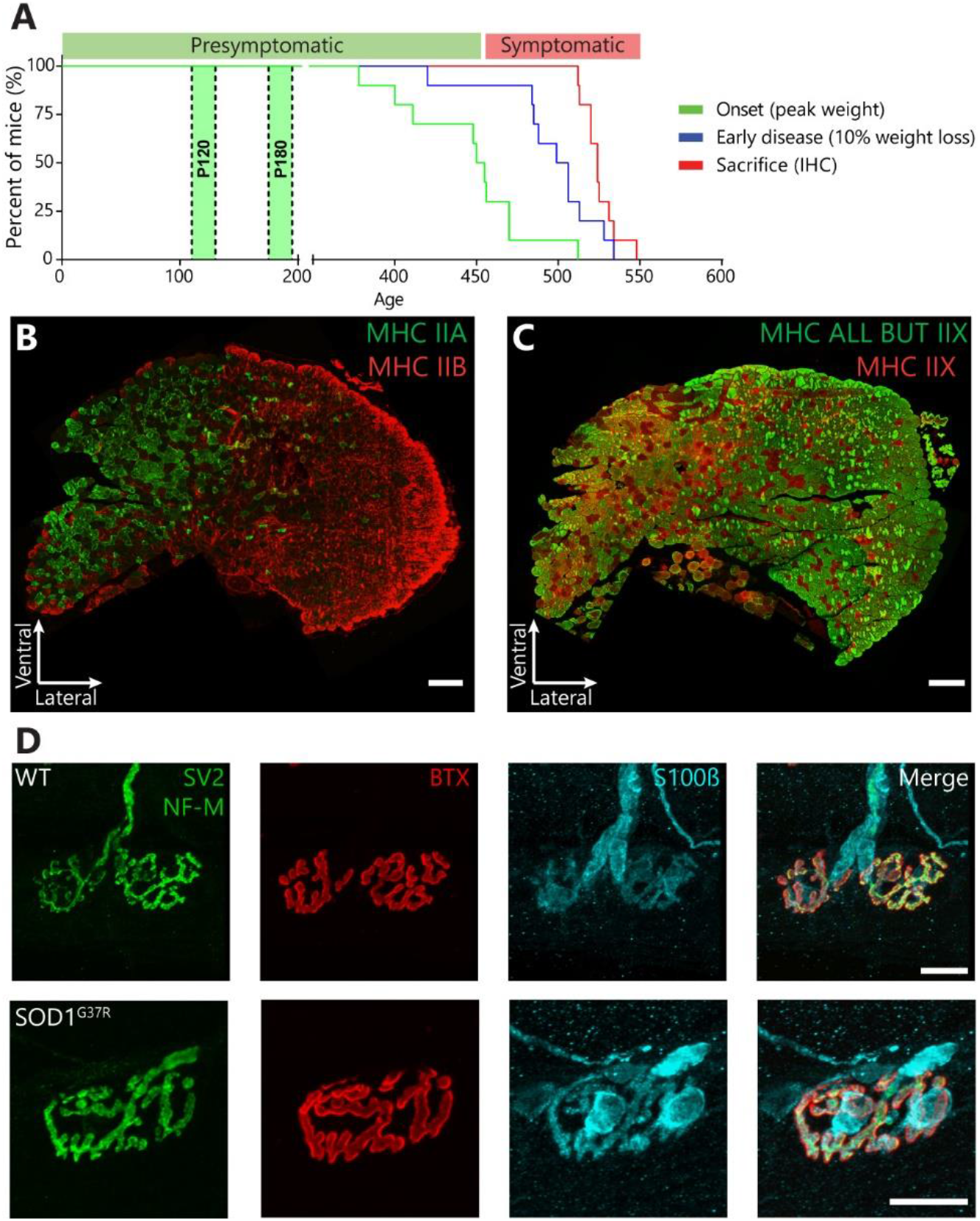
Disease progression in *SOD1*^*G37R*^ mice and fiber type composition of the *Sternomastoid* muscle. **(A)** Kalpan-Meier plot of the ages at which disease onset (green; median age: 456 days) and early disease (blue; median age: 506 days) are reached in *SOD1*^*G37R*^ mice under our conditions. All the symptomatic animals displayed similar motor phenotypes and were sacrificed (red, median 527 days) after the early disease stage. Experiments on presymptomatic animals were carried at P120 and P180. **(B-C)** Immunostaining of *Sternomastoid* cross-sections for Myosin Heavy Chain (MHC) type IIb (FF, red) and IIa (FR, green) (B) or all isotypes except IIx (all but IIx, green) and type IIx (FF, red) (C). Scale bars = 200 μm. (**D**) Representative examples of presynaptic nerve terminals (green, SV2 and NF-M), postsynaptic nAChR receptors red, (α-BTX) and glia (blue; S100β) labeling from the STM of P120 *WT* and *SOD1*^*G37R*^ mice. Scale bars = 20 μm.

### Phenotype evaluation

Starting at P290 or earlier, mice were weighted weekly to assess disease onset and progression as previously described (Lobsiger et al., 2009; Parone et al., 2013). Briefly, disease onset and early disease were defined as peak body weight and 10% loss from peak body weight respectively. Consistent with the recent reduction in transgene copy number in this line (Zwiegers et al., 2014), the median age of onset of the symptomatic animals used in this study was 456 days, while the median age of early disease was 506 days (Fig.1A; N=19). Phenotypically matched symptomatic animals were sacrificed for experimentation when 3 criteria were met (P505-P565) and are hereafter referred to as “symptomatic *SOD1*^*G37R*^ mice”: (1) they passed the early disease stage (10% weight loss); (2) they displayed hindlimb paralysis as assessed by tail suspension; and (3) they showed a reduced grid hanging time (less than 10 seconds). The average duration of the disease progression (from age of onset to sacrifice) was 78.26 ± 6.40 days.

### Sciatic nerve crush surgery

Mice were anesthetized using isoflurane (2-3% in 98-97% O_2_) in an induction chamber and anesthesia was then maintained under a breathing mask. Lubricant (Optixcare) was applied on the eyes to prevent dryness. Mice were placed in prone position and an incision was made on the left mid-thigh. The *gluteus maximus* and the *biceps femoris muscles* were delicately separated to expose the sciatic nerve. The nerve was then crushed with Moria microserrated curved forceps (MC31; maximal pressure for 15s). Muscles were then gently repositioned and the wound closed using 5-0 or 6-0 Vicryl suture (CDMV). Buprenorphine (3μg/10g of body weight; Temgesic) was subcutaneously administered three times during the following 24 hours. At the end of the surgery, mice were also injected subcutaneously with 0.25 mL of warm 0.9% saline twice to prevent dehydration. For sham-operated animals, the same procedure was followed, but the sciatic nerve was only exposed and not crushed.

### Sternomastoid calcium imaging

Both *STM* muscles and their nerves were dissected from the same animal in oxygenated Ringer REES solution containing (in mM): 110 NaCl, 5 KCl, 25 NaHCO_3_, 2 CaCl_2_, 1 MgCl_2_, 11 Glucose, 0.3 Glutamic Acid, 0.4 Glutamine, 5 BES (N,N-Bis(2-hydroxyethyl)taurine), 0.036 Choline Chloride, 4.34 × 10^−4^ Thiamine pyrophosphate (all Sigma-Aldrich). Muscles were pinned in a Sylgard184-coated (Dow Corning) recording chamber and the nerves were placed in two suction electrodes and were independently stimulated using a Pulsemaster A300 stimulator (square pulses: 20mV to 2V, 0.1ms duration; WPI). PSCs were loaded with the fluorescent Ca^2+^ indicator Rhod-3 (Life Technologies) by incubating muscles twice for 45 min in preoxygenated Ringer containing 5 μM Rhod-3 AM (Life Technologies), 0.02 % Pluronic acid (Life Technologies) and 0.15% dimethyl sulfoxide (DMSO) at 29 ± 2°C (Todd et al., 2010; Arbour et al., 2015). NMJs were located by labelling nAChRs prior to the start of the experiment with a closed-bath application of Alexa 488-conjugated α-BTX (5 μg for 10 min; Life Technologies). PSCs were easily identified using transmitted light microscopy and their identity was further confirmed through their apposition over BTX staining-. For experiments performed at P180, changes in fluorescence were monitored using a Zeiss LSM510 confocal microscope equipped with a 40x water-immersion lens (NA: 0.8; Zeiss). Rhod-3 was excited using a 543nm HeNe laser and emission was filtered using a 560nm long-pass filter. For experiments performed on symptomatic animals, changes in fluorescence were monitored using an Olympus FV-1000 confocal microscope equipped with a 60x water-immersion lens (NA: 0.9; Olympus). Rhod-3 was excited using a 559 nm diode laser and emission detected through a 570-625 nm spectral window. Changes in fluorescence were measured on the PSCs soma using the ImageJ software and were expressed as % ΔF/F_0_ = (F – F_0_)/F_0_. Recordings of PSC Ca^2+^-responses were not included when baseline fluorescence was unstable, focus changes occurred, spontaneous Ca^2+^ activity was observed or when Ca^2+^ levels did not return to the baseline after PSC activation. No direct comparisons between data obtained on different systems were made.

Ca^2+^ responses to endogenous neurotransmitter release were evoked by high-frequency stimulation of the nerve (50Hz, 20s). This pattern is known to elicit robust neurotransmitter release in this preparation (Wyatt and Balice-Gordon, 2008) and to efficiently trigger PSC Ca^2+^ responses in another fast-twitch muscle (Rousse et al., 2010). Alternatively, PSC mAChRs or purinergic receptors were activated by local application of agonists diluted in the extracellular Ringer REES solution, respectively muscarine (10 μM; Sigma) or ATP (1 μM; Sigma). Local drug application was achieved by applying a brief pulse of positive pressure (20 PSI, 150ms) using a Picospritzer II (Parker Instruments) through a glass micropipette (~2 μm tip, 5MΩ) positioned next to the NMJ. Because of the rundown of PSC Ca^2+^ responses following repeated activation (Jahromi et al., 1992; Rochon et al., 2001), local applications of agonists were performed before the high-frequency nerve stimulation was performed, each agonist was applied only once on each NMJ, and ATP and muscarine were applied at least 20 minutes apart on any given NMJ. Only a single high-frequency stimulation was executed on each muscle and PSC Ca^2+^-responses were imaged on a naïve NMJ. NMJs where high frequency nerve stimulation did not induce a presynaptic Ca^2+^ increase were discarded.

In all experiments, muscles were perfused (60-80 mL/h) with heated (28 ± 1 °C) Ringer REES containing 2.3 μg/mL of D-tubocurarine (Sigma) to prevent muscle contraction during nerve stimulation. In some experiments, mAChRs were blocked by adding the non-selective antagonist Atropine (10 μM; Sigma) to the extracellular Ringer REES solution. To ensure efficient blocking of mAChRs, muscles were perfused at least 45 min prior to the start of the experiment. PSCs responding to local application of muscarine after Atropine blockade were discarded.

### Whole-mount muscle preparations and immunohistochemistry

The *Sternomastoid* (STM) muscle is divided in two easily distinguishable parts on its ventral side: a lateral part composed exclusively of fast-fatigable (FF) motor-units (MUs) on the surface (the “white” part), and a medial part composed of a mixture of FF and fast-fatigue-resistant (FR) MUs (the “red” part) as inferred by their fiber type composition (Fig.1B et C) (Brichta et al., 1987). To exclusively evaluate vulnerable FF MUs in all experiments, we restricted our analysis to the “white” part of the STM which is easily observable using transmitted light. All surface NMJs analyzed in the *Extensor digitorum longus* (EDL) were associated with vulnerable FF MUs as previously shown (Tremblay et al., 2017). All NMJs analyzed in the *Soleus* (SOL) were associated with either fast-fatigue-resistant (FR) or slow (S) motor-units (MUs) as previously described (Pun et al., 2006; Arbour et al., 2015; Tremblay et al., 2017).

Immunohistochemical labeling was performed as described elsewhere (Todd et al., 2010; Darabid et al., 2013; Arbour et al., 2015). STM and SOL muscles were dissected in oxygenated (95% O_2_, 5% CO_2_) REES physiological solution and were pinned in a Sylgard-coated (Dow Corning) 10 mm petri dish and fixed for 10 min in 4% formaldehyde (Mecalab) diluted in PBS (in mM: 137 NaCl, 10 Na_2_HPO_4_, 2.7 KCl, 2 KH_2_PO_4_; all Sigma-Aldrich) at room temperature (RT). Then, muscles were permeabilized in 100% ice-cold methanol for 6 min at −20°C and non-specific labeling was reduced using 10 % normal donkey serum (NDS) diluted in PBS containing 0.01% Triton X-100 (20 min incubation). Muscles were incubated overnight at 4°C with a rat IgG2a anti-MAC-2 (Galectin-3) antibody (1:250; clone M3/38; Cedarlane, cat# CL8942AP), then 2h at RT with a rabbit IgG anti-S100β antibody (1:250; Dako-Agilent, cat# Z031101-2 or between 1:4 and 1:5, Dako-Agilent, cat#IR50461-2) followed by 2h at RT with the chicken IgY anti-Neurofilament medium chain (NF-M; 1:2000; Rockland Immunochemicals; cat# 212-901-D84) and the mouse IgG1 anti-synaptic vesicular protein 2 (SV2; 1:2000; Developmental Studies Hybridoma bank; concentrate) antibodies. Secondary antibodies donkey anti-rabbit IgG Alexa-594, donkey anti-rat IgG Alexa-647, goat anti-mouse IgG1 Alexa-488 and donkey anti-chicken Alexa-488 were incubated together for 1h at RT (all 1:500; Jackson Immunoresearch). All antibody dilutions were made in PBS buffer containing 0.01% Triton X-100 and 2% NDS. Finally, postsynaptic nicotinic receptors (nAChRs) were labeled using α-Bungarotoxin (BTX) conjugated with CF405 (4 μg/mL; Biotium; cat ##00002) for 45 min in PBS. In some cases (nerve crush experiment at P90, Fig.7A), the anti-SV2 antibody was replaced with goat anti-synaptotagmin (SyT; 1:250). In these cases goat anti-mouse IgG1 Alexa-405, donkey anti-chicken Alexa-405 (both 1:250; Jackson Immunoresearch), donkey anti-rabbit IgG Alexa 488 (1:500; Jackson Immunoresearch) and α-BTX conjugated to Alexa-594 (1.33 μg/mL; Life Technologies; cat# B13423) were used instead. After each incubation (except NDS blocking), muscles were rinsed three times for 5 min in PBS containing 0.01% Triton X-100. Muscles were mounted in ProlongGold anti-fade reagent (Life Technologies). Images were acquired on an Olympus FV1000 confocal microscope equipped with a 60x oil-immersion objective (N.A: 1.4; Olympus). Fluorescence was excited using a 488 nm (Alexa-488) argon laser and a 405 nm, a 559 nm and a 633 nm laser diode. Fluorescence emission was filtered using appropriate custom spectral windows (425 – 475nm, 500 – 545 nm, 570 – 630nm and 650 – 750nm). For figure representation, gamma was adjusted to enhance visibility of certain NMJ structures using Adobe Photoshop. No quantitative measurements were performed on the adjusted images.

For some experiments, muscles were fixed and immunolabeled after calcium imaging experiments to determine their innervations status post-hoc. In these experiments, confocal stacks of the BTX labeling pattern of the imaged and neighboring NMJs were acquired live and were used to re-identify the same NMJs following immunolabeling as previously described (Arbour et al., 2015).

### Gal-3 labeling on non-permeabilized whole-mount muscle tissue

To label extracellular *Gal-3*, muscles from Thy1-YFP animals were fixed in 4% formaldehyde and non-specific labeling was blocked using 10% NDS. Muscles were then incubated overnight at 4°C with the rat IgG2a anti-MAC-2 (Galectin-3) antibody (1:100), which was then revealed using a donkey anti-rat IgG Alexa-647 antibody (1:500; 60 min). Postsynaptic nAChRs were labeled with BTX conjugated to Alexa-594 (5μg/mL; 45 min). Muscles were rinsed three times for 5 min each between incubations and all dilutions and rinses were made in PBS buffer without Triton X-100. Muscles were then mounted in Prolong Diamond anti-fade reagent containing DAPI (Life technologies). Using this approach, no labeling of the presynaptic markers SV2 and NF-M could be observed, confirming that cells were not inadvertently permeabilized during the procedure (*data not shown*).

### Analysis of NMJ morphology

Analysis of NMJ morphology was performed using the seven criteria previously described and illustrated by Tremblay et al. (2017). Briefly, NMJ innervation and postsynaptic receptor organization (faint clustered or ectopic nAChRs) were analyzed and the presence of nerve terminal sprouting, polyinnervation and PSC process extensions was quantified. Although PSCs downregulate the S100β marker following denervation (Magill et al 2007), PSC somata on fully denervated endplates were still discernible, allowing quantification of Galectin-3 expressing PSCs. NMJs without any detectable S100β staining (presumably denervated for a long time) were not included in this study. Notwithstanding the limitations of S100β for the quantification of PSC process extensions in these conditions (Son and Thompson, 1995a), a large number of processes were labeled allowing quantification based on their presence on an NMJ rather than their number. Furthermore, we never observed a Galectin-3-positive S100β-negative process or a presynaptic sprout in absence of S100β staining, showing that S100β is reliable for this analysis in these conditions.

### Muscle cross-section immunohistochemistry

Immunostainning was performed similarly as previously described (Tremblay et al., 2017). Briefly, the *STM* muscle was dissected and frozen in cold optimal cutting medium compound (OCT; TissueTek) using isopentane at −80°C. Transverse cryosections (10 μm) were made and incubated in blocking solution (10% NDS in PBS). Sections were incubated with either mouse IgG1 anti-MHC type IIa (SC-71, 1:200), mouse IgG2b anti-MHC type I (BA-D5, 1:100) and mouse IgM anti-MHC type IIb (BF-F3, 1:200) or mouse IgM anti-MHC type IIx (6H1; 1:10) and mouse IgG1 anti-MHC all but IIx (BF-35, 1:200) for 1h at room temperature (all from Developmental Studies Hybridoma bank; concentrates (Schiaffino et al., 1989; Lucas et al., 2000)). They were revealed with appropriate secondary antibodies (1:500, Jackson Immunoresearch) and mounted in Prolong Gold antifade reagent (Life Technologies).

### Experimental design and statistical analyses

For Ca^2+^ imaging experiments, the number of animals used (N; biological replicates) and the number of PSCs analyzed (n; statistical replicates) are indicated in the text. Unpaired *t-tests* were performed to compare two different conditions from different experiments. *Two-way ANOVAs* tests were used to evaluate the effect of two independent variables on Ca^2+^-responses, using *Tukey’s multiple comparisons (Tukey’s test*) as a *post-test*. To analyze the heterogeneity of Ca^2+^ responses belonging to the same NMJs, the amplitude of the response of each cell was expressed as a percentage of the average response of all imaged cells on that NMJ. Then, the standard deviation of the resulting distribution for each group were compared using the *F test for unequal variance*, using *Holm-Sidak’s* method to correct for multiple comparisons. Statistical tests were performed in GraphPad Prism 7 software. Unless otherwise stated, all results are presented as mean ± SEM. All tests used a confidence interval of 95% (α = 0.05).

For the morphological analysis, all visible surface NMJs from left and right STM and SOL muscles were included. Number of animals used (N; representing both the biological and statistical replicates) and the number of NMJs or PSCs analyzed (n; number of observations) in each condition are indicated in the text. Due to the nature of the analysis (count data), the data does not meet the assumptions of following a Gaussian Distribution or having a variance equal between groups and symmetric (Crawley, 2007). Hence, *Generalized linear models (GLM)* were created, using a binomial error structure and a logit link function (logistic distribution), to analyze the results as previously described (Tremblay et al., 2017). For all criteria, the effect of the genotype (WT vs SOD1^G37R^) and the muscle (STM vs SOL) were evaluated. Furthermore, the effect of the innervation status of NMJs (fully innervated vs completely denervated) on PSCs’ repair response, identified by process extensions and Gal-3 expression, was also evaluated. The muscle and the innervation status were defined as within-subject factors. The effects of all *2-way* and *3-way* interactions were also evaluated. Finally, biologically relevant pairwise comparisons are illustrated in Figure 5A and were made using the estimated marginal means (EM means), using *Holm-Sidak’s* method to correct for multiple comparisons (hereafter referenced by *GLM post-test*). All analyses were made using the SPSS software (v.24.0.0.0; IBM). All tests used a confidence interval of 95% (α = 0.05).

## RESULTS

We have previously shown an increased activation of PSC mAChR during synaptic communication at S and FR NMJs in presymptomatic *SOD1*^*G37R*^ mice (P120 and P380) (Arbour et al., 2015). However, NMJs innervated by different MU-types display opposite changes in their synaptic properties in *SOD1*^*G37R*^ mice, i.e. neurotransmitter release is increased at S NMJs and decreased at FF NMJs (Tremblay et al., 2017). Knowing that alterations in synaptic activity can induce plastic changes in PSC activity (Belair et al., 2010), PSC excitability may be different at FF NMJs in *SOD1*^*G37R*^ mice. In the present study, we first tested whether PSCs at vulnerable FF NMJs also displayed an enhance mAChR activation during synaptic communication (mAChR contribution) in SOD1^G37R^ mice. For most experiments, the STM, (FF MUs on its ventro-lateral face; Fig. 1B and 1C), and the SOL (FR and S MUs; Arbour et al., 2015) nerve muscle preparations were used due to their similar rate of denervation in SOD1 mice despite their different MU type composition (Valdez et al., 2012). Their similar disease course allowed the analysis of the impact of the MU type on PSC repair properties under similar levels of endogenous denervation (symptomatic animals). FF NMJs in the STM of SOD1^G37R^ mice also showed a similar decrease in synaptic activity (*unpublished observations*) to FF NMJs in the EDL at P180 (Tremblay et al., 2017), indicating that, despite denervation occurring at later stages than in hindlimb fast-twitch muscles, early synaptic alterations are also present in the STM.

### Reduced PSC Ca^2+^ activity in the STM of presymptomatic SOD1^G37R^ mice

First, we examined the ability of PSCs to detect endogenous neurotransmitter release elicited by a motor nerve stimulation (50 Hz 20 sec) in the STM. PSC activity was monitored by imaging transient intracellular Ca^2+^ changes, a well-established reporter of their decoding activity (Rochon et al., 2001; Rousse et al., 2010). Experiments were performed at P120, age where the first alterations were observed in the SOL of *SOD1*^*G37R*^ mice (Arbour et al., 2015). At this stage, no signs of denervation were observed in the STM of *SOD1*^*G37R*^ mice (Fig. 1D; 50 NMJs, N=2).

At P120, all PSCs of *WT* (18 PSCs; 9 NMJs; N=6) and *SOD1*^*G37R*^ mice (19 PSCs; 7 NMJs; N=4) responded to nerve stimulation-induced neurotransmitter release in the STM. PSC Ca^2+^ response amplitude was similar between *WT* and *SOD1*^*G37R*^ mice (333.7 ± 31.1 % ΔF/F_0_, n=18 vs 279.2 ± 27.4 % ΔF/F_0,_ n=19 respectively; t_(35)_=1.320; *p*=0.1954 *Unpaired t-test*). This is consistent with the different progression of limb and trunk muscles and the observation that PSC Ca^2+^ responses were unchanged in the SOL 2 months earlier (P60) (Arbour et al., 2015).

We next examined PSC properties in the STM at P180 to test whether changes in PSC activity were delayed in the STM compared to the SOL. Again, PSCs responsiveness to nerve stimulation was similar between *SOD1*^*G37R*^ and *WT* mice (*SOD1*^*G37R*^: 31 out of 33 PSCs; 12 NMJs; N=8; *WT*: 20 out 22 PSCs; 11 NMJs; N=8). However, unlike at P120, PSC Ca^2+^ responses at P180 were significantly smaller in *SOD1*^*G37R*^ mice (Fig 2A and 2D; *WT-Ctrl* : 403.2 ± 22.8 % ΔF/F_0_, n=20 vs *SOD1-Ctrl*: 270.6 ± 24.3 % ΔF/F_0_, n=31; Effect of genotype: F_(1,77)_=25.13; *p<0.*0001, *Two-way ANOVA; p*=0.0005, *Tukey’s test on Two-way ANOVA*). Hence, PSC properties were also altered in the STM of *SOD1*^*G37R*^ mice at an early presymptomatic stage, but in an opposite manner to what was observed in the SOL (Arbour et al., 2015).

**Figure 2:**
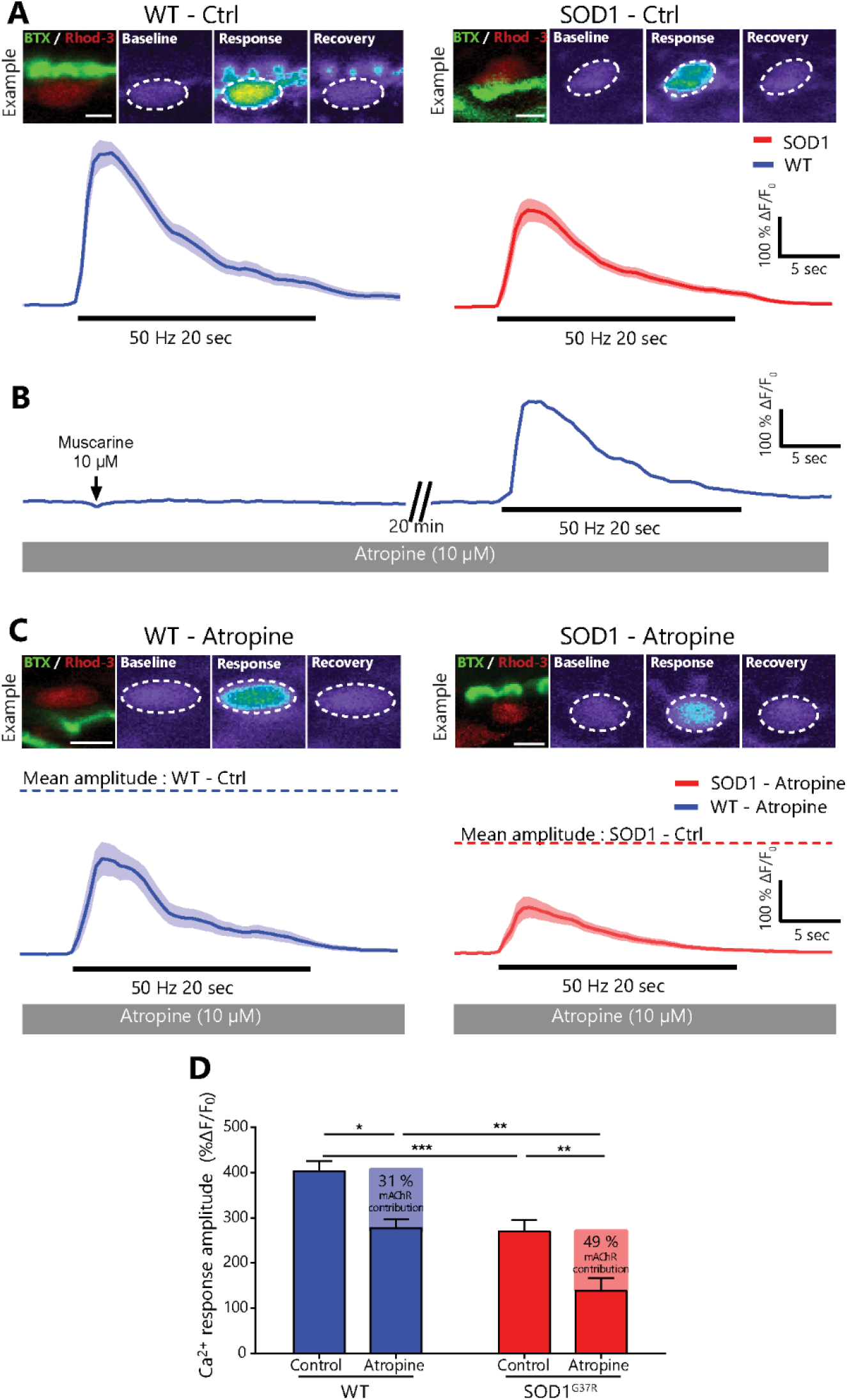
Relative increase of mAChR-dependent PSC activity in *SOD1*^*G37R*^ mice at a presymptomatic stage. **(A)** Average PSC Ca^2+^ responses ± SEM (pale area) in *SOD1*^*G37R*^ mice (*right)* and *WT* controls (*left*) induced by high-frequency nerve stimulation at P180 in absence of atropine (Ctrl). *Top*, localization of the PSC on an endplate (left, apposed over BTX staining) and representative false color images of the changes in Rhod-3 fluorescence representing changes in intracellular Ca^2+^ levels before, during and after motor nerve stimulation. **(B)** Example of PSC Ca^2+^ responses in *WT* controls evoked by local application of muscarine (*left*) or nerve stimulation (*right*) in presence of 10 μM atropine. Note the blockade of muscarine-induced Ca^2+^ response and the reduced amplitude of nerve stimulation-evoked Ca^2+^ responses (l*eft*, compared with **A**). **(C)** Average PSC Ca^2+^ responses ± SEM (pale area) in *SOD1*^*G37R*^ mice (*right)* and *WT* controls (*left*) induced by high-frequency nerve stimulation at P180 in presence of 10 μM atropine. Note the reduced amplitude of nerve stimulation-evoked Ca^2+^ responses compared with the average peak amplitude without atropine (dashed lines) *Top*, localization of the PSC on an endplate (left) and representative false color images of the changes in Rhod-3 fluorescence representing changes in Ca^2+^ levels before, during and after motor nerve stimulation. **(D)** Histogram of the amplitude of PSC Ca^2+^ responses in all conditions P180. Scale bar = 5 μm. *, *p* <0.05; **, *p*<0.01; ***, *p*<0.005.

### Relative increase of mAChR-dependent PSC activity in STM at a presymptomatic stage

Next, we tested whether the contribution of mAChR activation to PSC Ca^2+^ activity (mAChR contribution) was increased in the STM, despite the smaller PSC Ca^2+^ responses.

For this, Ca^2+^ imaging was performed in presence of the general mAChR antagonist Atropine (10 μM) at P180. As shown in Figure 2B, blockade of PSC mAChRs was confirmed before the nerve stimulation by the inability of local muscarine applications (10 μM) to elicit Ca^2+^ responses. Out of the 32 cells analyzed (14 in *WTs* and 18 in *SOD1s*), only 2 PSCs from *WT* mice exhibited a small response to local application of muscarine in presence of atropine and were discarded. In the presence of atropine, all PSCs responded to nerve stimulation-induced neurotransmitter release in both *WT* (12 PSCs; 7 NMJs; N=4) and *SOD1*^*G37R*^ mice (18 PSCs; 8 NMJs; N=4). As expected, the amplitude of PSC Ca^2+^ responses was significantly decreased in both groups (Effect of atropine: F_(1,77)_=22.63 *p*<0.0001; *Two-way ANOVA;* Interaction: F_(1,77)_=0.01446, *p=*0.9046). In *WT* mice, it was reduced from 403.2 ± 22.8 % ΔF/F_0_, (n=20) to 277.5 ± 18.4 % ΔF/F_0_ in presence of atropine (n=12; *p*=0.0194; *Tukey’s test on Two-way ANOVA*). This represents a 31% contribution of mAChRs to PSC Ca^2+^ responses that is similar to our observation in the adult SOL (Arbour et al., 2015). Likewise, PSC Ca^2+^ responses in *SOD1*^*G37R*^ mice had a significantly smaller amplitude in presence of atropine (Fig. 2D; *SOD1-* Ctrl: 270.6 ± 24.3 % ΔF/F_0_, n=31 vs *SOD1*-Atropine: 138.5 ± 27.6 % ΔF/F_0_, n=18; *p*=0.0013; *Tukey’s test on Two-way ANOVA*). However, this represents a 49% mAChR contribution in *SOD1*^*G37R*^ mice, which is 1.6-fold larger than in *WT* animals. Hence, the contribution of mAChR activation to PSC Ca^2+^-activity is also increased in the STM of *SOD1*^*G37R*^ mice.

Furthermore, PSC Ca^2+^ responses in *SOD1*^*G37R*^ mice remained significantly smaller than in *WT* mice in presence of atropine (*p*=0.0094; *Tukey’s test on Two-way ANOVA*). As such, the reduction of the amplitude of PSC Ca^2+^ responses in the STM of *SOD1*^*G37R*^ seem to be attributable to the reduced activation of non-muscarinic PSC receptors. The other major type of receptors by which PSCs detect synaptic activity at the mammalian NMJ are purinergic receptors (Rochon et al., 2001). However, their specific contribution to nerve-evoked PSC Ca^2+^-response was not evaluated as the underlying set of receptors remain undetermined at the adult mammalian NMJs (Rochon et al., 2001; Arbour et al., 2015).

As a whole, these results suggest that a relative increase in the contribution of mAChR to PSC activity is a common pathological feature between both fast-twitch and slow-twitch muscles in *SOD1*^*G37R*^ mice.

### Heterogeneous nerve-induced PSC activity at individual NMJs in the STM at a presymptomatic stage

Interestingly, PSCs belonging to the same NMJ in the STM of *SOD1*^*G37R*^ mice displayed heterogeneous responses to nerve stimulation, an unseen characteristic in *WT* mice (Fig. 3A). When only considering NMJs where multiple PSCs were imaged, PSC Ca^2+^ responses were all within ± 20 % of the average response on their NMJ in *WT* animals (Fig. 3B; SD = 11.17 %; N=8; n=20). However, in *SOD1*^*G37R*^ mice, PSC Ca^2+^ responses ranged from −99 % up to +85 % of the average response on their NMJ (Fig. 3B; SD = 43.65 %; N=8; n=31), which is significantly more than in *WT* mice (F_(30,19)_=15.27; *p*= 0.0004; *F-test*). This suggests an uneven alteration of PSCs excitability at presymptomatic stage.

**Figure 3:**
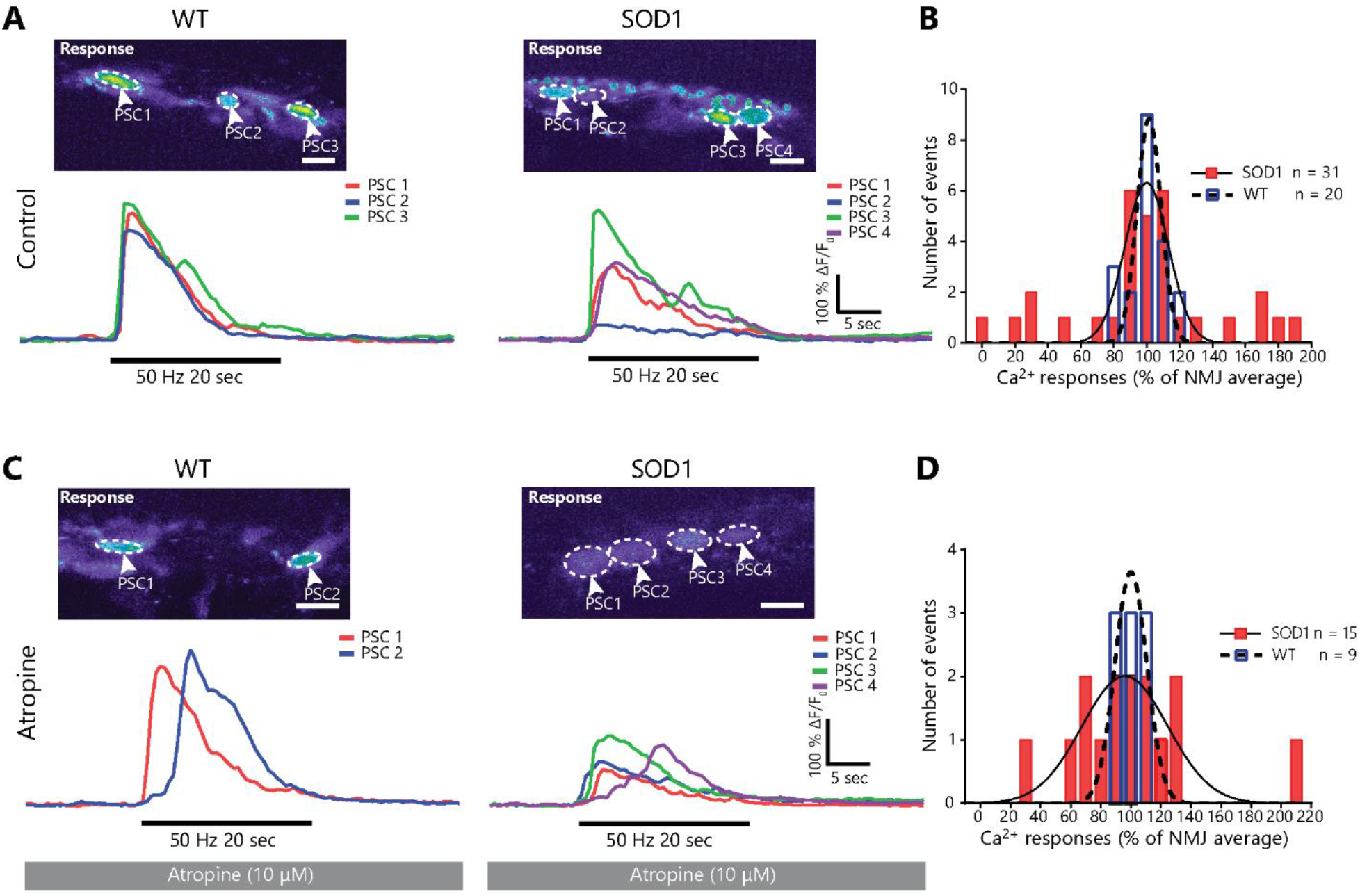
Muscarinic receptor-independent heterogeneity in PSC activity at individual NMJs in the STM of presymptomatic *SOD1*^*G37R*^ animals. **(A)** Examples of PSC Ca^2+^ responses from a single NMJ in *SOD1*^*G37R*^ mice (*right*) and *WT* controls (*left*) in absence of Atropine. Note the greater heterogeneity in Ca^2+^ responses in *SOD1*^*G37R*^ mice. **(B)** Frequency distribution of PSC Ca^2+^ in *WT* (blue, dashed line) and *SOD1*^*G37R*^ (red, solid line) mice normalized to the average response of all PSCs on their NMJ. **(C)** Example of individual PSC Ca^2+^ responses from an NMJ in *SOD1*^*G37R*^ mice (*right*) and *WT* controls (*left*) in presence of Atropine. **(D)** Frequency distribution of PSC Ca^2+^ in *WT* (blue, dashed line) and *SOD1*^*G37R*^ (red, solid line) mice normalized to the average response of all PSCs on their NMJ in presence of atropine. Lines in **B** and **C** represent best-fit Gaussian curves for each sample (*WT*: dashed line; *SOD1*^*G37R*^: solid line) and are used for representation purposes only. Scale bar = 10 μm.

These results could imply that the increased mAChR-dependent PSC activity is also uneven in the STM of *SOD1*^*G37R*^ mice and therefore may not be a common feature to all PSCs. If this were the case, blockade of PSC mAChRs would restore homogeneity in Ca^2+^ responses over a single NMJ. However, heterogeneity in PSC Ca^2+^ responses was maintained following blockade of mAChRs in *SOD1*^*G37R*^ mice (Fig. 3C). In presence of atropine, PSC Ca^2+^ responses in *WT* animals varied between −20 % to +30 % of each other on a given NMJ (Fig. 3D; N=4; n=9; SD=15.12 %;) while PSC Ca^2+^ responses in *SOD1*^*G37R*^ mice varied between −71 % and +113 % (Fig. 3D; N=3; n=15; SD=47.51 %), thus maintaining the difference observed in control condition (F_(14,8)_=9.877; *p*=0.0087; *F-test*). Indeed, presence of atropine did not significantly affect the variability of Ca^2+^-responses on single NMJs in *WT* (*WT*-Ctrl vs *WT*-Atropine: F_(8,19)_=1.832; *p*=0.4610; *F-test*) or in *SOD1*^*G37R*^ mice (*SOD1*-Ctrl vs *SOD1*-Atropine: F_(14,30)_=1.185; *p*=0.6702; *F-test*). These results reveal that heterogeneity in PSC Ca^2+^ responses in *SOD1*^*G37R*^ mice is independent of mAChR activation. Hence, increased relative mAChR contribution to PSC synaptic decoding appears to be present in all PSCs in the STM of *SOD1*^*G37R*^ mice.

### Reduced PSC detection of purinergic signals in STM of presymptomatic SOD1^G37R^ mice

PSC Ca^2+^ activity heavily depends on their intrinsic properties (Belair et al., 2010; Rousse et al., 2010; Darabid et al., 2013; Arbour et al., 2015), such as the presence and sensitivity of their muscarinic and purinergic receptors. To further investigate PSC properties in the STM of SOD1^G37R^ mice, we first tested PSCs sensitivity to cholinergic signals by monitoring PSC Ca^2+^ responses elicited by local application of the mAChR agonist muscarine (10 μM). As shown in Figure 4A, PSCs in both groups exhibited similar responsiveness (*WT:* 28 out 32; 17 NMJs; N=10 vs *SOD1:* 32 out of 33; 16 NMJs; N=8) and Ca^2+^ responses of similar amplitude following muscarine application (Fig. 4B; 191.6 ± 23.1 % ΔF/F_0_, n=28 vs 228.0 ± 23.0 % ΔF/F_0_, n=32, respectively; t_(58)_=1.127; *p*=0.2644; *Unpaired t-test*). Thus, these results are consistent with the suggested reduced activation of another type of PSC receptors during synaptic communication in the STM of *SOD1*^*G37R*^ mice.

**Figure 4:**
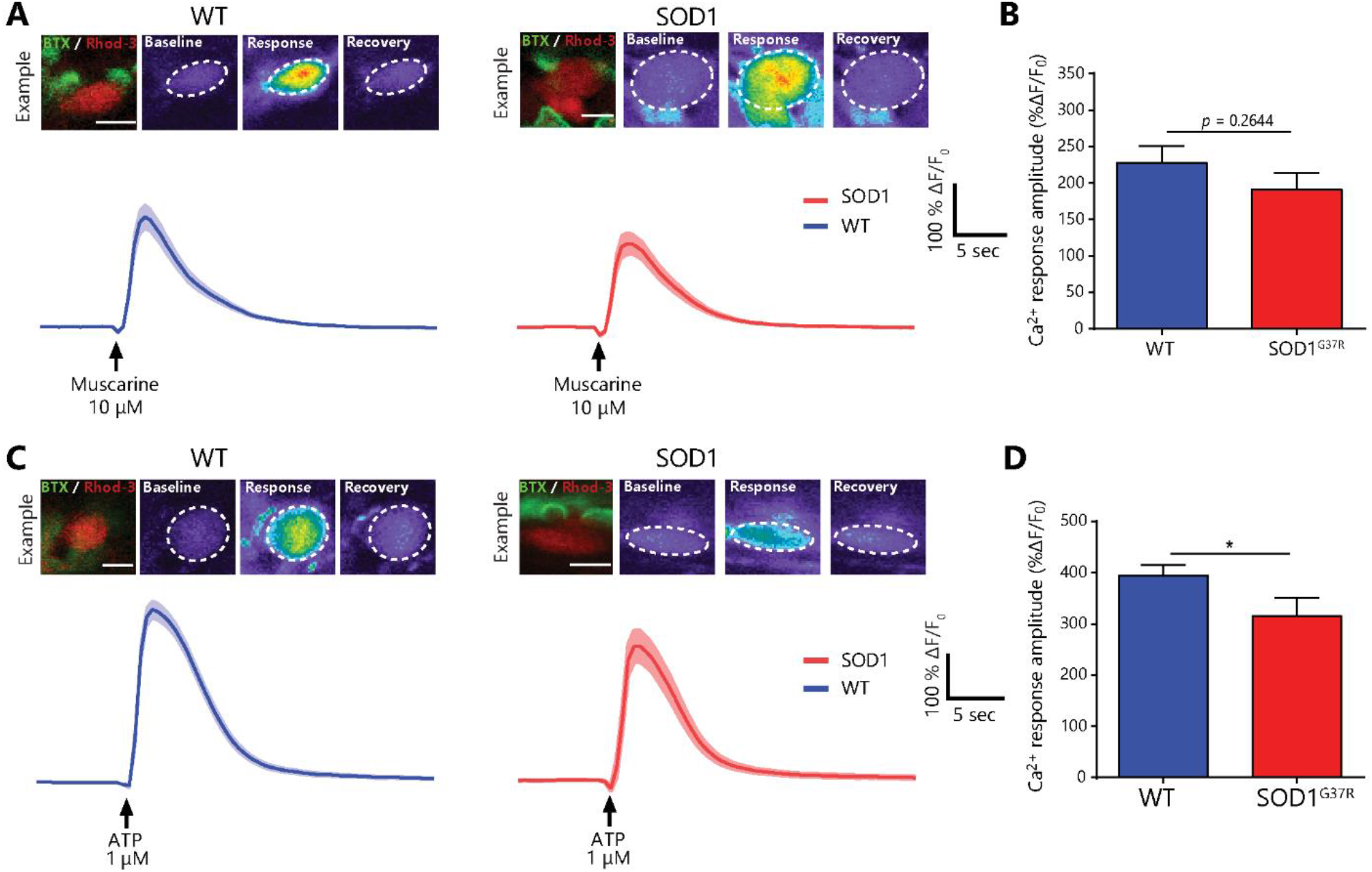
PSC Ca^2+^ responses elicited by local application of muscarine or ATP in presymptomatic *SOD1*^*G37R*^ mice. **(A)** Average PSC Ca^2+^ responses ± SEM (pale area) in *SOD1*^*G37R*^ mice (*right)* and *WT* controls (*left*) induced by local application of muscarine at P180. *Top*, localization of the PSC on an endplate (left, apposed over BTX staining) and representative false color images of the changes in Rhod-3 fluorescence representing changes in intracellular Ca^2+^ levels before, during and after the response. **(B)** Histogram of the amplitude of PSC Ca^2+^ responses induced by muscarine application at P180. **(C)** Average PSC Ca^2+^ responses ± SEM (pale area) in *SOD1*^*G37R*^ mice (*right)* and *WT* controls (*left*) induced by local application of ATP at P180. *Top*, localization of the PSC on an endplate and representative false color images of the changes in Rhod-3 fluorescence representing changes in Ca^2+^ levels before, during and after the response. **(D)** Histogram of the amplitude of PSC Ca^2+^ responses induced by ATP application at P180. Scale bar = 5 μm. *, *p* <0.05.

Next, PSCs’ sensitivity to purinergic agonists was examined by monitoring PSC Ca^2+^ responses after local application of the general purinergic agonist ATP (1μM). Local application of ATP robustly triggered a Ca^2+^ response in 100% of PSC in both groups (*WT:* 32 out of 32; 15 NMJs; N=9 vs *SOD1:* 21 out 21; 13 NMJs; N=7). However, as shown in Figure 4C, ATP-induced PSC Ca^2+^ responses in the STM of *SOD1*^*G37R*^ mice were smaller than in *WT* mice (Fig. 4D; 315.2 ± 35.6 % ΔF/F_0_, n=21 vs 394.3 ± 20.2 % ΔF/F_0_, n=32, respectively; t_(51)_=2.078; *p*=0.0429; *Unpaired t-test*). These results suggest that PSCs are less sensitive to purinergic signals in the STM of *SOD1*^*G37R*^ mice.

Altogether, our results obtained by imaging PSC Ca^2+^ activity show that the relative contribution of mAChR to PSC activity is increased, while PSCs’ sensitivity to purinergic signals is decreased at FF NMJs in SOD1^G37R^ mice. Despite having slightly different properties depending on the MU type they are associated with (Arbour et al., 2015), these results show that the contribution of mAChR to PSCs’ activity increases at all NMJ types in SOD1^G37R^ mice.

### PSCs express and secrete the phagocytic marker Gal-3 following denervation in WT mice

Knowing that PSC mAChR activation regulates gene expression and represses their repair phenotype, we postulated that altered muscarinic properties would be associated with defects in PSC-dependent NMJ repair mechanisms. Notably, PSCs become phagocytic and participate in the clearance of nerve terminal debris following denervation (Duregotti et al., 2015) similarly to axonal SCs (Reichert et al., 1994; Brosius Lutz et al., 2017). Axonal debris clearance is essential for efficient NMJ reinnervation, cellular debris hindering axonal growth in the endoneural tube (Kang and Lichtman, 2013).

Interestingly, phagocytic axonal SCs express the β-galactoside-binding lectin Galectine-3 (Gal-3, a.k.a MAC-2) (Reichert et al., 1994; Rotshenker et al., 2008), similarly to other phagocytic glial cells (Rotshenker, 2009; Nguyen et al., 2011; Morizawa et al., 2017), suggesting that its expression could also reflect PSC phagocytic activity. However, Gal-3 expression has never been evaluated in PSCs. Thus, we first investigated whether PSCs expressed Gal-3 at denervated NMJs in *WT* animals by performing an IHC for the three synaptic components (presynaptic, postsynaptic and glia) and Gal-3. We induced the complete denervation of SOL NMJs in adult *WT* animals by sciatic nerve crush and monitored the presence of Gal-3 two days later (Fig. 5A; Nerve crush: N=3, n=43; Sham: N=3, n=42). As expected, we observed a robust expression of Gal-3 in the soma and processes of PSCs following sciatic nerve crush (Fig. 5A, E; 85.66 ± 2.02 % of PSCs, n=134) compared to sham surgery (2.883 ± 1.92 % of PSCs, n=105; *p*<0.001; *GLM post-test*). These results confirm that PSCs express Gal-3 following NMJ denervation as part of their normal response to injury in *WT* animals.

**Figure 5:**
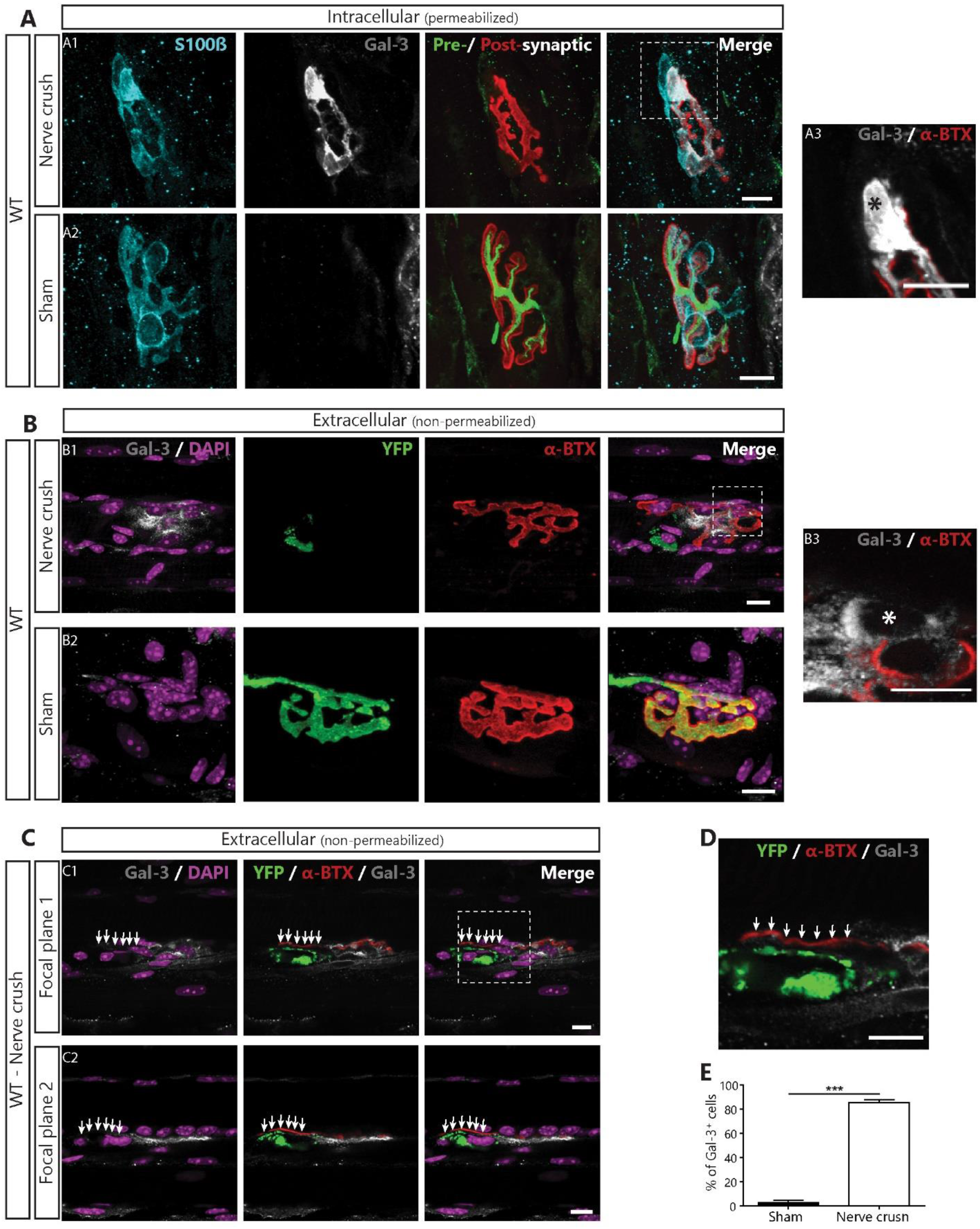
PSCs express the phagocytic marker Gal-3 following denervation. **(A)** Representative confocal images of immunolabeling of glial cells (blue; S100β), Gal-3 (white), presynaptic nerve terminals and postsynaptic nAChR (merged: respectively green, SyT; red, α-BTX,) from the SOL of *WT* mice following a sciatic nerve crush (A1), or in Sham operated animal (A2). (A3) High magnification image of the region identified in A1 showing that Gal-3 is present inside the soma (*). **(B)** Representative images of the extracellular labelling of postsynaptic nAChR (red), Gal-3 (white) and nuclei (DAPI; purple) from the SOL of Thy1-YFP mice following sciatic nerve crush (B1) or Sham surgery (B2). (B3) High magnification image of the region identified in B1 showing that Gal-3 is present around the soma (*). Note that Gal-3 appears to be both intracellular and extracellular following NMJ denervation. **(C)** Two single confocal planes of a denervated NMJ showing that YFP^+^-debris accumulation on the endplate (arrows) correlates with the lack of extracellular Gal-3 following sciatic nerve crush. **(D)** High magnification of the region in C1 showing that Gal-3 is absent next to YFP^+^-debris. **(E)** Percentage of PSCs expressing Gal-3. Quantification is based on permeabilized immunolabeling. Scale bar = 10μm. ***, p<0.005.

Gal-3 can be present intracellularly, secreted or associated with transmembrane proteins (Dumic et al., 2006; Yang et al., 2008). Previous studies on axonal SCs suggested that Gal-3 may exert its function in the extracellular space (Reichert et al., 1994). To further characterize its role at the NMJ, we examined whether Gal-3 was present in the extracellular space two days after sciatic nerve crush by performing an IHC in absence of membrane permeabilization. Since our presynaptic markers are intracellular, the efficacy of the denervation was validated by performing experiments on animals expressing the “yellow fluorescent protein” (YFP) in all motor neurons (homozygous Thy1-YFP16 mice). As shown in Figure 5B, Gal-3 was present in the extracellular space following sciatic nerve crush (Fig. 5B_1_; N=3, n=45) but not sham surgery (Fig. 5B_2_; N=2, n=27). Extracellular Gal-3 seemed membrane bound as it was present all around PSCs’ somata (Fig. 5B_3_). However, close comparison of Gal-3 staining in permeabilized and non-permeabilized tissue (Fig 5A_3_ and 5B_3_ respectively) reveals that part of the staining was missing in non-permeabilized tissues (asterisk). This difference shows that Gal-3 was also present in the cytoplasm and possibly in the nucleus. Interestingly, accumulation of YFP^+^ axonal debris on the endplate often correlated with the lack of extracellular Gal-3 on neighboring PSCs (Fig. 5C and D, arrows; 6 out of 7 observations). Altogether, these results show that Gal-3 is expressed by PSCs following denervation, is present in the extracellular space at the NMJ and correlates with efficient clearance of presynaptic debris on the endplate.

### Some PSCs fail to upregulate Gal-3 following denervation in presymptomatic SOD1^G37R^ mice

Based on our Ca^2+^-imaging results, we hypothesized that altered PSCs’ decoding properties would be associated with an inadequate phagocytic phenotype on denervated NMJs in presymptomatic *SOD1*^G37R^ mice. Thus, we tested if PSCs upregulated Gal-3 following an experimentally-induced denervation in presymptomatic SOD1^G37R^ mice (Sciatic nerve crush at ~P180). We evaluated PSCs Gal-3 expression in the EDL (FF NMJs on its surface, (Tremblay et al., 2017)) and the SOL (FR and S NMJs) to assess any MU-type dependent differences. The EDL was used instead of the STM for nerve-crush experiments, since the denervation of SOL and EDL NMJs can be induced using the same surgical procedure (sciatic nerve crush), thereby reducing inter-animal and inter-procedure variability. Furthermore, the surgery to induce the denervation of STM NMJs is much more invasive, causing a strong inflammatory response in the muscle (data not shown) and possibly altering PSC properties. Based on our hypothesis, we predict that PSC Gal-3 expression would be reduced at denervated NMJs of both types in SOD1^G37R^ mice.

Interestingly, fewer PSCs upregulated Gal-3 in the EDL and the SOL of P180 SOD1^G37R^ mice than of littermate controls two days after sciatic nerve crush (Figure 6A and B, arrows; Figure 6C; WT EDL: N=4, 83 NMJs, n=246 PSCs; WT SOL: N=4, 108 NMJs, n=288 PSCs; SOD1 EDL: N=4, 90 NMJs, n=252 PSCs; SOD1 SOL: N=4, 89 NMJs, 205 PSCs, Effect of genotype: *p* < 0.001, *GLM*). Importantly, PSC Gal-3 expression was also dependent on the muscle type (Figure 6C, Effect of muscle: *p* < 0.001, *GLM*; Interaction between genotype and muscle: *p* = 0.028, GLM) such that significantly fewer PSCs expressed Gal-3 following denervation in the EDL than in the SOL in WT mice (Figure 6C, 74.65 ± 4.73% vs 88.25 ± 1.65% respectively, *p* = 0.022, *GLM post-test*) and in SOD1^G37R^ mice (Figure 6C, 42.00 ± 3.85% vs 81.25 ± 3.01% respectively; *p* < 0.001, *GLM post-test*). Even though more PSCs failed to upregulate Gal-3 in both muscles in SOD1^G37R^ mice compared to WT mice (Figure 6C, EDL: WT vs SOD1, *p* < 0.001; SOL: WT vs SOD1 *p* = 0.022, *GLM post-test*), this effect was much more pronounced in the EDL than in the SOL. Hence, PSCs seem less likely to adopt a phagocytic phenotype at denervated NMJs in presymptomatic SOD1^G37R^ mice in a manner consistent with NMJ vulnerability in ALS. These results suggest that PSCs may not adequately promote NMJ repair upon the endogenous denervation of NMJs during the symptomatic phase in *SOD1*^*G37R*^ mice.

**Figure 6:**
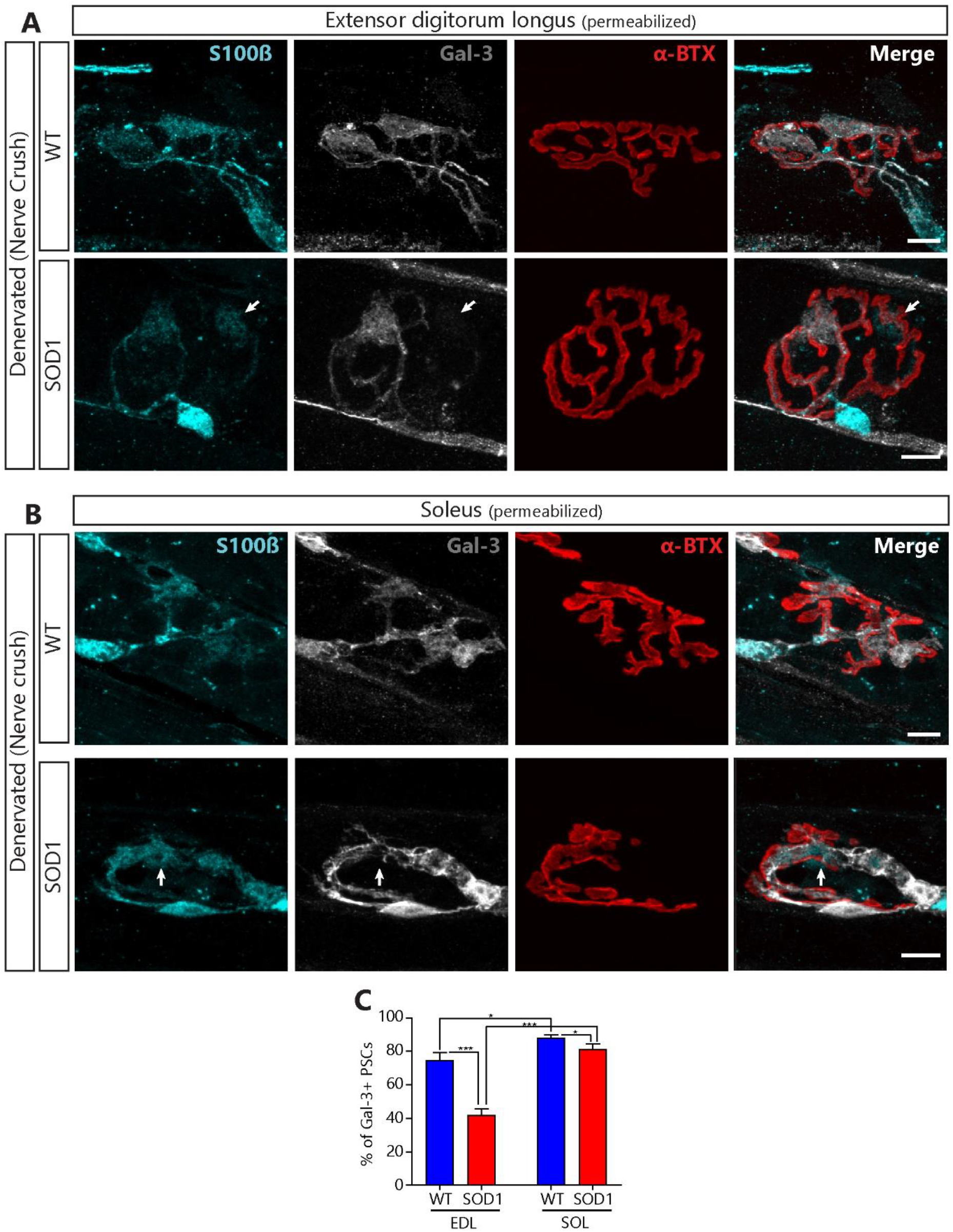
Lower expression of Gal-3 in PSCs of presymptomatic *SOD1*^*G37R*^ mice following denervation. Representative confocal images of the immunolabeling of glial cells (blue; S100β), Gal-3 (white) and postsynaptic nAChR (red, α-BTX,) from the EDL (**A**) or the SOL (**B**) of *WT* (top) and *SOD1*^*G37R*^ mice (bottom) two days after a sciatic nerve crush. Note the absence of Gal-3 expression in some of the PSCs from SOD1 mice (arrows). **(C)** Quantification of the percentage of PSCs expressing Gal-3 two days following a sciatic nerve crush as a function of the genotype and the muscle type. Scale bar = 10μm. *, *p* <0.05; ***, *p*<0.001.

### PSC Ca^2+^ activity is no longer altered in the STM of symptomatic SOD1^G37R^ mice

We previously showed that PSC muscarinic receptor activation remained elevated until the pre-onset stage (P380) in the SOL (Arbour et al., 2015). Therefore, we asked if the alteration of PSCs properties at FF NMJs would also persist until the symptomatic stage. To address this question, we first examined PSC Ca^2+^ activity in the STM of phenotypically matched late symptomatic SOD1^G37R^ mice and age-matched *WT* mice. PSC Ca^2+^ activity was induced by high-frequency nerve stimulation (50Hz 20 sec) or by local application of muscarine (10 μm) or ATP (1μm).

Among PSCs that met our inclusion criteria (see material and methods), responsiveness to nerve stimulation, local application of muscarine or local application of ATP was similar between groups (Nerve stimulation: *SOD1*^*G37R*^ 12 out of 12 PSCs, N=5 vs *WT* 13 out of 13 PSCs N=6; Muscarine: *SOD1*^*G37R*^ 43 out of 52 PSCs, N=8 vs *WT* 33 out of 39 PSCs, N=6; ATP: *SOD1*^*G37R*^ 49 out of 49 PSCs, N=8 vs *WT* 39 out of 39 PSCs, N=6). Although some PSC Ca^2+^ responses to endogenous transmitter release in *SOD1*^*G37R*^ mice tended to be slightly smaller than in *WT* mice, the difference was not statistically significant (Figure 7A-B; 294.8 ± 35.7 % ΔF/F_0_, n=12 vs 353.4 ± 31.2 % ΔF/F_0,_ n=13 respectively; t_(23)_=1.227; *p*=0.2324 *Unpaired t-test*).

**Figure 7:**
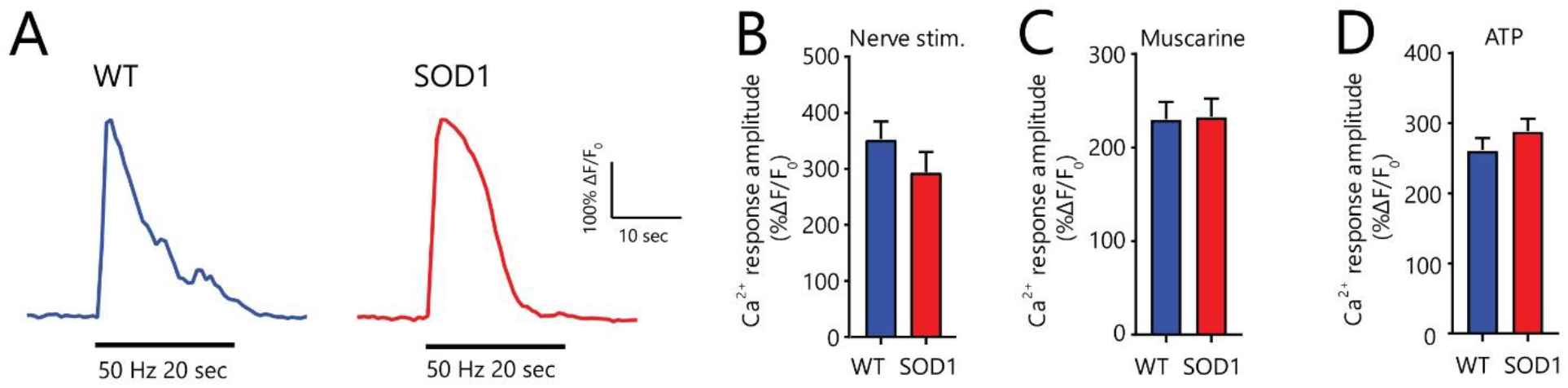
Unchanged PSC properties in phenotypically-matched symptomatic *SOD1*^*-G37R*^ mice. (**A**) Representative examples of PSC Ca^2+^ responses induced by high-frequency nerve stimulation in *WT* (blue) and *SOD1*^*G37R*^ (red) mice. **(B-D)** Histograms of the amplitude of PSC Ca^2+^ responses induced by high-frequency nerve stimulation (**B**) and local application of 10μM muscarine (**C**) or 1 μM ATP (**D**).

Similarly, the amplitude of PSC Ca^2+^ responses to local application of muscarine and ATP were similar between *WT* and symptomatic *SOD1*^*G37R*^ mice (Figure 7C-D; Muscarine : 230.6 ± 18.8 % ΔF/F_0_, n=33 vs 233.3 ± 19.1 % ΔF/F_0,_ n=43 respectively; t_(74)_=1.007; *p*=0.9200 *Unpaired t-test*; ATP : 261.8 ± 16.9 % ΔF/F_0_, n=39 vs 288.9 ± 17.2 % ΔF/F_0,_ n=49 respectively; t_(86)_=1.106; *p*=0.2718 *Unpaired t-test)*. Hence, PSC Ca^2+^ properties seem to have evolved during disease progression in the STM, with Ca^2+^ responses to endogenous transmitter release or local ATP and muscarine applications no longer being reduced in symptomatic *SOD1*^*G37R*^ mice.

### Similar level of denervation in STM and SOL muscles of symptomatic SOD1^G37R^ mice

We next wanted to evaluate if PSC’s ability to upregulate Gal-3 or to promote NMJ repair was still impaired in symptomatic *SOD1*^*G37R*^ mice. Using IHC, PSCs’ repair properties (Gal-3 expression, PSC process extensions and nerve terminal sprouting) were evaluated at all NMJ types under similar levels of denervation, by comparing the STM and the SOL of phenotypically matched late symptomatic *SOD1*^*G37R*^ animals (P505-565; Fig. 1A). This experimental design allowed us to isolate the effect of the muscle type on the repair properties since, unlike the STM, denervation levels in leg fast-twitch muscles, such as the EDL, are drastically higher than in the SOL at this stage, with only few NMJs being fully innervated (*unpublished observations* and Tremblay et al. 2017). Observations were analyzed as a function of the innervation state of the NMJs (innervated vs denervated) in addition to the muscle type (STM vs SOL) and the genotype (*WT* vs *SOD1*^*G37R*^) as illustrated in Figure 8A.

**Figure 8:**
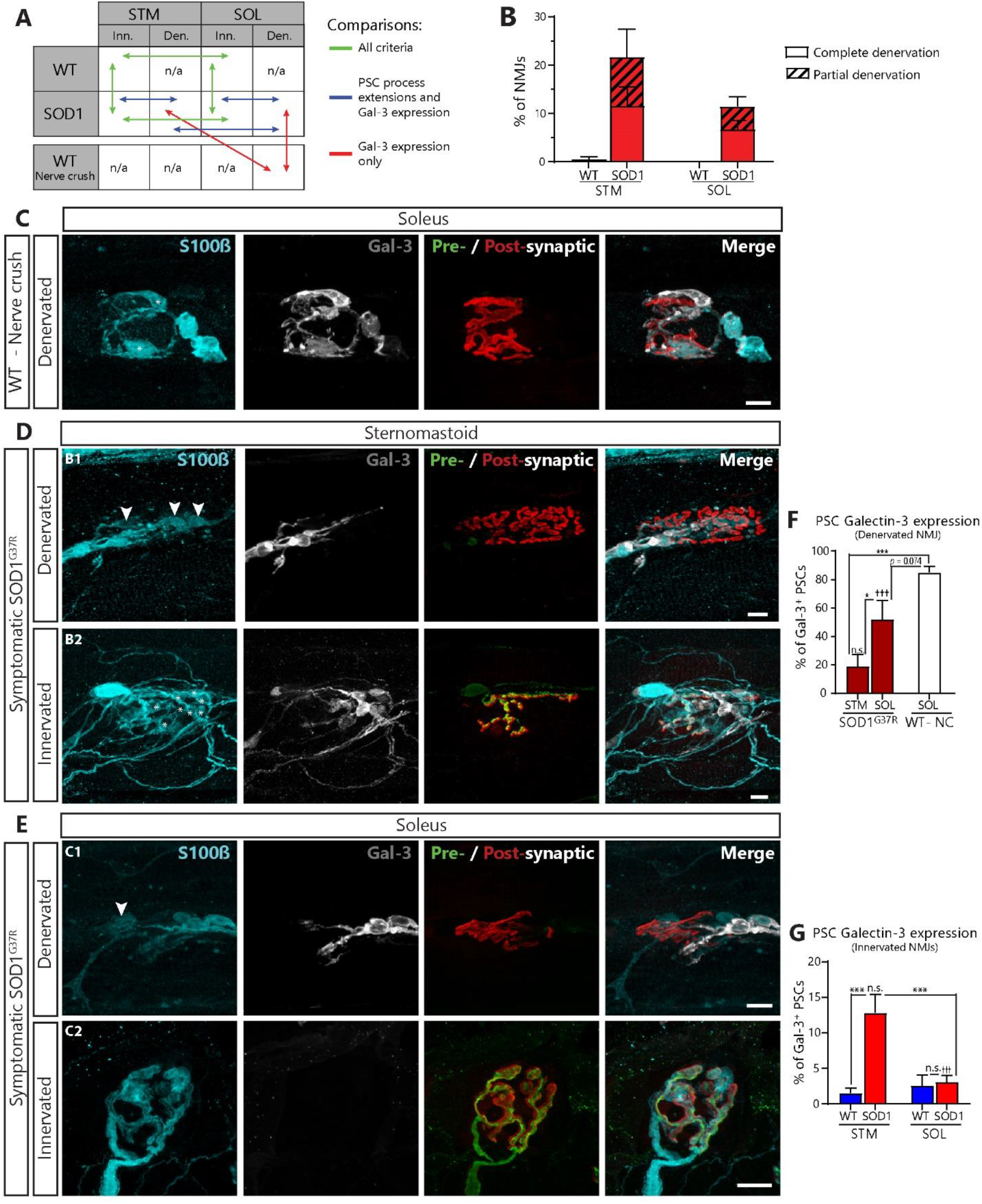
Paradoxical expression of Gal-3 in the STM and SOL of symptomatic *SOD1*^*G37R*^ mice. **(A)** Schematic of the biologically relevant pairwise comparisons to assess the effect of genotype (*WT* vs *SOD1*^*G37R*^), muscle type (STM vs SOL) and innervation status (innervated vs denervated). **(B)** Histogram of the percentage of completely and partially denervated NMJs across all groups. Den., denervated; Inn., innervated. (**C-E**) Immunolabeling of glia (blue; S100β), Gal-3 (white), presynaptic nerve terminals and postsynaptic nAChR receptors (merged: respectively green, SV2 and NF-M; red, α-BTX). **(C)** Representative confocal images of NMJs from the SOL of aged *WT* mice following nerve-crush induced denervation. **(D)** Representative confocal images of denervated (B1) and innervated (B2) NMJs from the STM of symptomatic *SOD1*^*G37R*^ mice. Note the presence of Gal-3 in PSCs (*) on an innervated NMJ in the STM of *SOD1*^*G37R*^ mice. Also note the disorganization of PSC processes. **(E)** Representative confocal images of denervated (C1) and innervated (C2) NMJs from the SOL of *SOD1*^*G37R*^ mice. Note the presence of Gal-3 in axonal Schwann cells and its absence from PSCs (B and C, arrowheads). **(F)** Percentage of PSCs expressing Gal-3 on denervated NMJs and, **(G)** innervated NMJs in all groups. Symbols on the top of histogram bars represent the difference between innervated and denervated NMJs (D vs E). Scale bars = 10 μm. ns, nonsignificant; ***, *p*<0.005; †††, *p*<0.005.

First, we needed to confirm that the level of NMJ denervation in each muscle were comparable. Consistent with previous observations in the *SOD1*^*G93A*^ mice (Schaefer et al., 2005; Valdez et al., 2012), we confirmed that the extent of denervation (partial or complete) was comparable between the STM and the SOL in symptomatic *SOD1*^*G37R*^ mice, not being significantly higher in the STM than in the SOL (Fig. 8B; STM: 21.71 ± 5.76%, N=9 vs SOL: 11.51 ± 1.98%, N=7; *p*=0.116; *GLM post-test*). Indeed, we found that 11.63 ± 3.91% of NMJs in the STM were completely denervated (31 out of 266, N=9) compared to 6.80 ± 1.84%; in the SOL (20 out of 282, N=7). Similar results were obtained for partially denervated NMJs (STM, 10.08 ± 3.36%; 29 out of 266 NMJs, N=9 vs SOL, 4.11 ± 1.97%; 15 out of 282, N=7). Of note, only 1 out of the 156 NMJs analyzed in the STM of *WT* animals (0.54 ± 0.54%; N=6) was completely denervated and none were partially denervated. In the SOL, none of the 152 NMJs analyzed in *WT* animals (N=5) showed any sign of denervation. A partially denervated NMJ could be either undergoing denervation or re-innervation, which are two very distinct physiological states (Martineau et al., 2018). Due to this uncertainty, partially denervated NMJs were excluded from the following analyses.

As intended with this experimental design, these observations confirm that a possible difference in the level of denervation will not bias the analysis of Gal-3 expression or PSC repair properties at symptomatic stage.

### Lack of Gal-3 at denervated NMJs in symptomatic SOD1 animals

We examined Gal-3 expression in PSCs NMJs in the STM and the SOL of symptomatic SOD1^G37R^ mice. To rule out age-related changes in the debris clearance pathway (Kang and Lichtman, 2013), we also performed a sciatic nerve crush in aged *WT* mice (P450-P500). PSCs in the SOL of aged *WT* mice robustly expressed Gal-3 following sciatic nerve crush (Fig. 8C; 85.03 ± 4.07 %, N=4, n=120) and at similar levels than in P180 mice (Figure 5). Hence, PSCs maintain their ability to upregulate Gal-3 in 14-16 month-old WT mice.

As expected, we found that PSC Gal-3 expression varied according to the state of innervation and the muscle in *SOD1*^*G37R*^ mice and in age-matched controls (Effect of genotype: *p*<0.001, Effect of innervation: *p*<0.001, Genotype*innervation interaction: *p*<0.001; Muscle*innervation interaction: *p*<0.001, *GLM*). Interestingly however, numerous PSCs did not express Gal-3 on completely denervated NMJs in *SOD1*^*G37R*^ mice in both muscles (Fig. 8D_1_ and 8E_1_, arrowheads). Consistent with our finding in presymptomatic mice (Fig. 6), significantly less PSCs expressed Gal-3 in the STM of *SOD1*^*G37R*^ animals when compared to nerve crush-induced denervated NMJs in *WT* mice (Fig. 8F; 18.94 ± 8.32 % of PSCs; n=136 vs 85.03 ± 4.07 % of PSCs n=120; *p*<0.001; *GLM post-test*). Furthermore, presence of Gal-3 was highly variable in the SOL, ranging from 0.00% to 77,78% of PSCs (51.81 ± 13.56 %, n=49), and an overall trend towards lower levels was observed when compared to nerve crush-induced denervation (Fig. 8F; SOD1^G37R^ SOL vs Nerve crush WT SOL; *p*=0.074; *GLM post-test*). Interestingly, PSCs Gal-3 expression on denervated NMJs was significantly higher in the SOL than in the STM (Fig. 8F; SOD1^G37R^ SOL vs STM; *p*=0.020; *GLM post-test*), again reflecting the higher vulnerability of FF NMJs. Importantly, axonal SCs in both muscles robustly expressed Gal-3 when associated with denervated NMJs, suggesting that this alteration is specific to PSCs (Fig 8D_1_ and E_1_). Altogether these results show that PSCs unreliably upregulate Gal-3 on denervated NMJs in symptomatic *SOD1*^*G37R*^ mice, especially on vulnerable NMJs.

### Gal-3 is expressed on some innervated and partially innervated NMJs in the STM of symptomatic SOD1 mice

Paradoxically, while Gal-3 was absent from PSCs on denervated STM NMJs in *SOD1*^*G37R*^ mice, it was expressed in some PSCs on fully innervated NMJs in the STM (Fig. 8D_2_, asterisks and 8G; *SOD1*^*G37R*^: 12.84 ± 2.54 % of PSCs, n=866 vs *WT*: 1.55 ± 1.68 % of PSCs, n=618; *p*<0.001; *GLM post-test*). More importantly, a similar proportion of PSCs expressed Gal-3 whether the NMJ was innervated or denervated in the STM of *SOD1*^*G37R*^ mice (Fig. 8F-G, 12.84 ± 2.54 % vs 18.94 ± 8.32 %; *p*=0.132; *GLM post-test*) suggesting that Gal-3 expression in PSCs was independent of the state of innervation in this ALS model. This upregulation of Gal-3 on innervated NMJs was not observed in the SOL of *SOD1*^*G37R*^ animals (Fig. 8E_2_; *SOD1*^*G37R*^ SOL Innervated: 3.12 ± 0.92%, n=708 vs *WT* SOL Innervated: 2.59 ± 1.50, n=435; *p*=0.231; *GLM post-test*), where its expression was specific to denervated NMJs (Innervated: 3.12 ± 0.92% vs Denervated : 51.81 ± 13.56 %; *p*<0.001; *GLM post-test*). Gal-3 expression was also not observed at innervated NMJs in presymptomatic (P120-180) and pre-onset (P350) *SOD1*^*G37R*^ mice (data not shown).

In line with this paradoxical observation, a number PSCs on partially innervated NMJs also expressed Gal-3 and were usually associated with the nerve terminal in the STM of symptomatic *SOD1*^*G37R*^ mice (Fig. 9A_1_). Again, this was not observed in the SOL (Fig. 9A_2_). Indeed, 20 out of 29 (69.97 %) partially innervated NMJs had at least 1 Gal-3^+^-PSC in the STM compared to only 2 out 15 (13.33 %) in the SOL. In 80% of these cases in the STM Gal-3^+^-PSCs were preferentially associated with the presynaptic nerve terminal. Altogether, these results are suggestive of active PSC phagocytosis of presynaptic elements on denervating or reinnervating NMJs.

**Figure 9:**
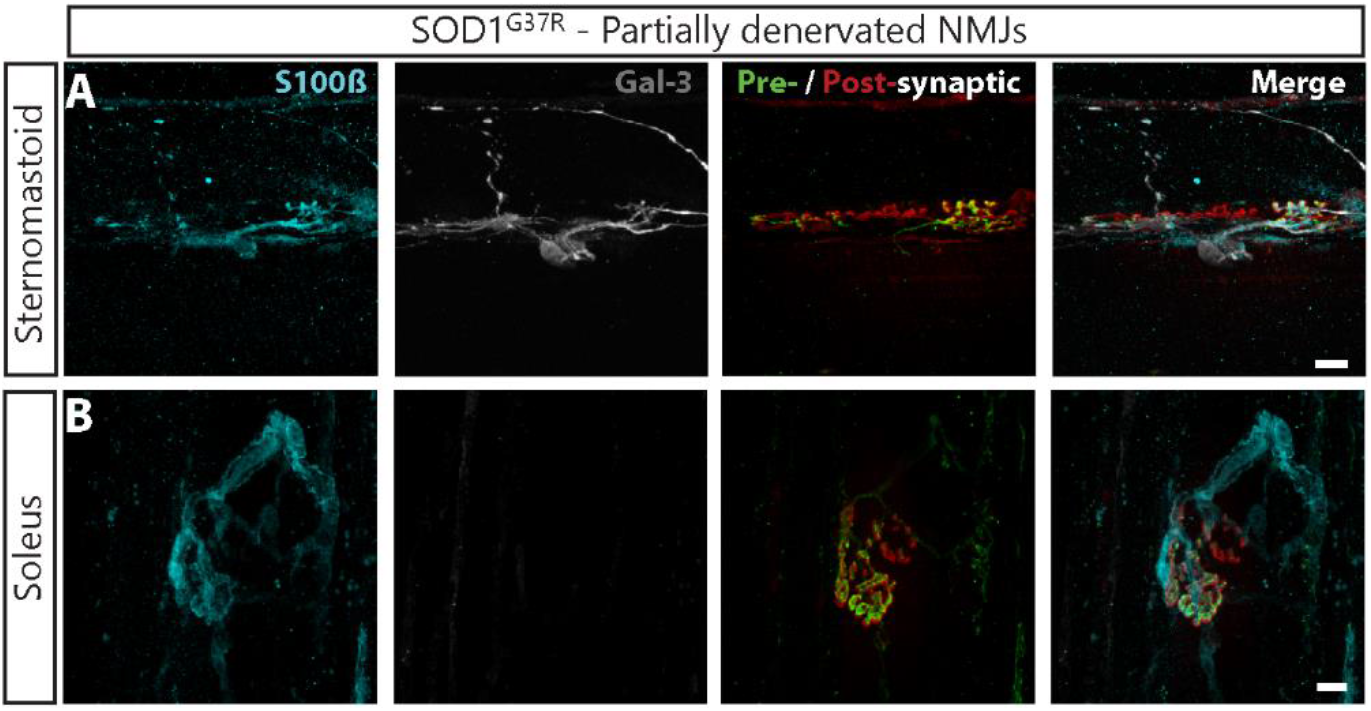
Gal-3 in PSCs associated with nerve terminals on partially innervated NMJs in the STM of symptomatic *SOD1*^*G37R*^ mice. Immunolabeling of glia (blue; S100β), Gal-3 (white), presynaptic nerve terminals and postsynaptic nAChR receptors (merged: respectively green, SV2 and NF-M; red, α-BTX). **(A)** Representative confocal images of partially innervated NMJs from the STM and **(B)** the SOL of symptomatic *SOD1*^*G37R*^ mice. Note that Gal-3^+^-PSCs are preferentially associated with the innervated part of the endplate rather than the denervated part. Scale bars = 10 μm

### PSCs extend disorganized processes in both the STM and the SOL

Next, we characterized another fundamental PSC mechanism contributing to NMJ repair: the extension of glial processes and the guidance of terminal sprouting. Importantly, this aspect of PSC repair properties could not be evaluated in presymptomatic animals following a complete nerve crush, since this procedure only poorly induces the extension of PSC processes (Love and Thompson, 1999). We monitored the presence and targeting of PSC process extensions in symptomatic *SOD1*^*G37R*^ animals. These extensions are normally observed from denervated NMJs towards nearby innervated NMJs, a phenomenon essential in the initiation and guidance of nerve terminal sprouting (Reynolds and Woolf, 1992; Son and Thompson, 1995b, a; Son et al., 1996; O’Malley et al., 1999).

In general, we found significantly more NMJs associated with extended PSC processes in SOD1^G37R^ than in WT mice (Fig. 10A and B; Effect of genotype: *p*<0.001, *GLM*) but this effect varied slightly according to the muscle type (Interaction genotype*muscle: *p*=0.044; *GLM*). Denervated NMJs were more likely to be associated with extended PSC processes than innervated NMJs (Effect of innervation: *p*<0.001, *GLM*), but this effect also varied depending on the muscle (Interaction innervation*muscle: *p*=0.013, *GLM*).

**Figure 10:**
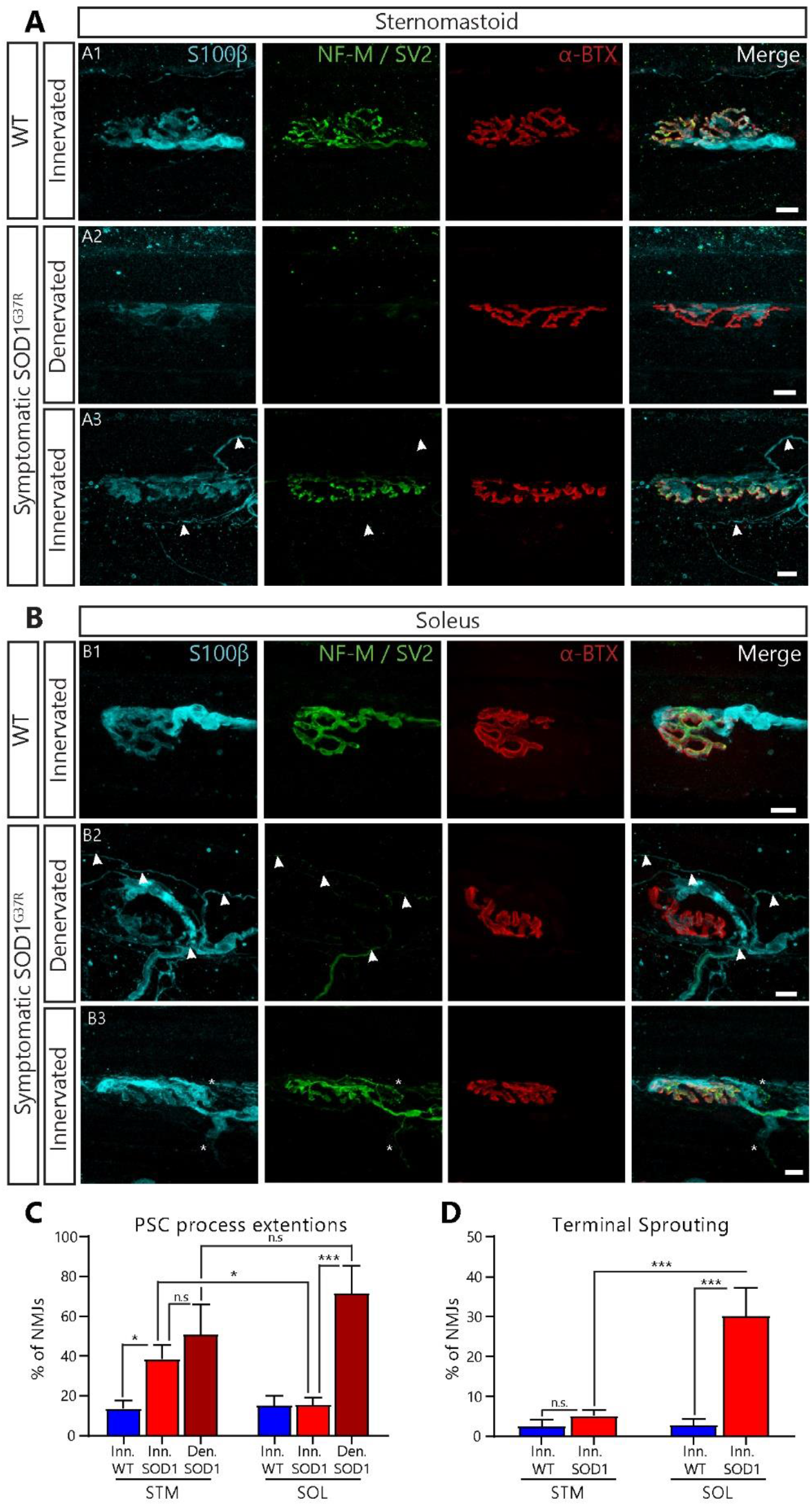
Abnormal and disorganized nerve terminal sprouting and PSC process extension in the STM and the SOL of symptomatic *SOD1*^*G37R*^ mice. Immunolabeling of glial cells (blue; S100β), presynaptic nerve terminals (green; NF-M and SV2) and postsynaptic nAChR receptors (red; α-BTX). **(A)** Representative confocal images of NMJs from the STM of age-matched *WT* controls (A1) and of symptomatic *SOD1*^*G37R*^ mice (A2: denervated; A3: innervated). **(B)** Representative confocal images of NMJs from the SOL of *WT* animals (B1) and of symptomatic *SOD1*^*G37R*^ mice (B2: denervated; B3: innervated). Note the nerve terminal sprouting “avoiding” the nearby denervated NMJ (B2; arrowheads). **(C)** The percentage of NMJs associated with PSC processes in the STM or the SOL according to the state of innervation. **(D)** Percentage of innervated NMJs displaying nerve terminal sprouting. Scale bars = 10 μm. ns, nonsignificant; *, *p* <0.05; ***, *p*<0.005.

Indeed, in the STM, completely denervated NMJs were not significantly more likely to be associated with extended PSC processes than innervated NMJs in *SOD1*^*G37R*^ mice (Fig. 10A_3_ arrowheads and C; 51.05 ± 14.99%, n=31 vs 38.48 ± 7.11%, n=206, respectively; *p*=0.858; *GLM post-test*). Conversely, innervated NMJs in the STM of *SOD1*^*G37R*^ mice were more likely to be associated with extended glial processes than innervated NMJs in *WT* mice (Fig. 10A and C, 13.73 ± 4.05% n=155; *p*=0.047; *GLM post-test*). Hence, PSCs in the STM of *SOD1*^*G37R*^ extended their processes but this did not seem to be dependent on the innervation state of NMJs. These results contrast with those obtained in the SOL. Indeed, PSCs on denervated NMJs in the SOL of SOD1^G37R^ mice robustly extended their processes compared to PSCs on innervated NMJs (Fig. 10B and C, 71.67 ± 13.76, n=36 vs 15.65 ± 3.68, n=247; *p*<0.001; *GLM post-test*). Hence, while PSC process extensions from denervated NMJs occurred as often in both muscles (*p*=0.858; *GLM post-test*), they were abnormally more likely to be associated with innervated NMJs in the STM than in the SOL (*p=*0.047; *GLM post-test*).

Importantly though, these results also show that 49 % of denervated NMJs in the STM and 28% in the SOL lacked PSC process extensions (see Fig.6A_2_ for example). These proportions are higher than the expected 1% based on studies using partial denervation (Son and Thompson, 1995b) and, in most cases, only one process per NMJ was observed (1.13 ± 0.67 processes on average) (Son and Thompson, 1995b; O’Malley et al., 1999). Furthermore, PSC processes were often highly disorganized in the STM and SOL of SOD1^G37R^ mice (47.09 ± 9.87 % and 31.02 ± 18.62 % of processes on average respectively; see Fig 10A_3_, 10B_2_ and 10B_2_ for example), often changing direction, not following muscle fibers and frequently making loops around themselves. Hence, although present, PSC process extensions were often disorganized and less abundant than expected on denervated NMJs regardless of the muscle type. These results further suggest that PSC properties are heterogeneous, which is reminiscent of the heterogeneity in PSC decoding properties observed at a presymptomatic stage.

### Nerve terminal sprouting is absent in the STM and disorganized in the SOL

Extended PSC processes contacting nearby innervated NMJs can initiate and guide nerve terminal sprouting back to denervated NMJs (Son and Thompson, 1995b; Son et al., 1996; O’Malley et al., 1999). However, previous studies reported conflicting results whereby nerve terminal sprouting is absent (Schaefer et al., 2005; Tallon et al., 2016) or abundant in *SOD1*^*G93A*^ mice (Valdez et al., 2012). Hence, we next tested whether nerve terminal sprouting would be present, albeit disorganized, on innervated NMJs in *SOD1* animals. Based on the lower, but not null, ability of FF NMJs to elaborate terminal sprouts (O’Malley et al., 1999; Frey et al., 2000; Wright et al., 2009), we expected fewer terminals sprouts in the STM compared with the SOL.

In general, we found significantly more NMJs associated with nerve terminal sprouts in the SOL of SOD1^G37R^ animals than any other groups (Fig. 10A - C; Effect of genotype: *p*=0.001; Effect of muscle: *p*<0.001; Interaction genotype*muscle: *p*=0.015; *GLM*). Indeed, 30.26±6.96% of innervated NMJs in the SOL of *SOD1*^*G37R*^ mice (Fig. 10B_3_; n=259) formed sprouts compared to 2.89 ± 1.46% of SOL NMJs in *WT* mice and 5.26 ± 1.37% of STM NMJs in *SOD1*^*G37R*^ mice (Fig. 10A_3_, 10B_1_ and 10D; n=152 and n=206, *p*<0.001 and *p*<0.001 respectively; *GLM post-test*) which suggests an active re-innervation effort in the SOL. Consistent with previous observations made on FF NMJs though (Martineau et al., 2018), we did not observe significantly more nerve terminal sprouts at STM NMJs in *SOD1*^*G37R*^ mice than in *WT* mice (5.26 ± 1.37%, n=206 vs 2.56 ± 1.64%, n=155; *p*=0.311). This finding was nevertheless surprising considering that over a third of them were being contacted by extended PSC processes (Fig.10A_3_ for example). Therefore, initiation of nerve terminal sprouting in symptomatic *SOD1* mice appears deficient in the STM, but not the SOL, resulting in “empty” extended PSCs processes contacting both innervated and denervated endplates (Fig. 10C).

Importantly though, nerve terminal sprouts in the SOL were often disorganized, going in many directions and even “avoiding” nearby denervated NMJs (Fig. 10B_2_, arrowheads). Indeed, 33.46 ± 6.50 % of sprouting NMJs had disorganized sprouts in the SOL of *SOD1*^*G37R*^ mice, which is consistent with the muddled PSC process extensions. As a whole, these results show that nerve terminal sprouting was almost completely absent in the STM and disorganized in the SOL of symptomatic *SOD1*^*G37R*^ animals.

In summary, these results show that PSCs display a paradoxical Gal-3 expression in symptomatic *SOD1*^*G37R*^ animals, being inconsistent on denervated NMJs of both muscles and unexpectedly present on innervated and partially innervated NMJs in the STM. This suggests that PSCs do not adopt a phagocytic phenotype upon denervation of vulnerable NMJs, and also to a lesser extent of partially resistant NMJs. Along with the disorganized processes and the abnormal nerve terminal sprouting, these results suggest that PSCs do not adopt an appropriate phenotype for re-innervation in ALS. Importantly, these main findings were replicated in the STM of a small number of pre-onset (P70) SOD1^G93A^ animals. Namely, PSCs on denervated NMJs failed to upregulate Gal-3 and to extend processes, resulting in the absence of nerve terminal sprouting (*data not shown*). Altogether, these results show that alteration PSC repair properties are not specific to a single *SOD1* model.

## DISCUSSION

Our results reveal inappropriate glial repair properties at the NMJs of a fast- and a slow-twitch muscle in symptomatic *SOD1* animals. These changes are associated with an altered PSC excitability, long before disease onset. Despite multiple differences in PSC properties (decoding and repair) between muscles and the heterogeneity in PSC properties, our data converges towards the same end results whereby PSCs on denervated NMJs inconsistently adopted a phagocytic phenotype, formed muddled processes, failed to guide nerve terminal sprouting appropriately and presented an increased mAChR-dependent activity, albeit transiently in a fast twitch muscle. Thus, PSC excitability and repair properties are incompatible with NMJ reinnervation across all MU types in *SOD1* models.

### NMJ repair defects in ALS

Here, we show that numerous PSC-dependent NMJ repair mechanisms are abnormal in *SOD1* animals, suggesting that NMJ reinnervation may be deficient during the progression of ALS. Although compensatory reinnervation is known to occur during the disease (Schaefer et al., 2005; Martineau et al., 2018), this process may be suboptimal, resulting in inefficient or unstable NMJ repair. Such bungled compensation could contribute to the loss of motor function in ALS by exacerbating loss of NMJs. Consistent with this hypothesis, numerous studies documented delays in NMJ reinnervation following axonal injury in ALS models (Sharp et al., 2005; Pun et al., 2006; Henriques et al., 2011; Swarup et al., 2012; Mesnard et al., 2013; Carrasco et al., 2016b). These observations suggest that a number of core NMJ repair mechanisms are altered in these models. For instance, an intrinsically slower axonal growth or cellular debris accumulation in the endoneural tube and at the NMJ that would deter NMJ reformation after denervation (Kang and Lichtman, 2013). Our observation that PSCs fail to adopt a phagocytic phenotype on denervated NMJs (Fig. 8) is consistent with these mechanisms. However, since the inadequate Gal-3 expression was specific to PSCs, and was not observed for axonal SCs, our results suggest that other mechanisms besides debris accumulation in the endoneural tube could contribute to the delay in NMJ reinnervation after nerve injury.

Importantly, global nerve injuries do not recapitulate the progressive denervation occurring in ALS. Hence, evaluating NMJ repair following partial denervation would be more instructive as it would allow compensatory reinnervation to take place through nodal (axonal) and nerve terminal sprouting of intact motor axons (Thompson and Jansen, 1977; Son and Thompson, 1995b). Since PSCs unreliably extended muddled processes (Fig. 10), we predict that reinnervation based on these sprouting events would also be drastically impaired. Indeed, despite being more abundant than in controls, PSC processes were less abundant than expected (Son and Thompson, 1995b; O’Malley et al., 1999) which could underlie the reduced terminal sprouting observed following blockade of neurotransmission in *SOD1* mice (Frey et al., 2000).

Interestingly, Schaefer et al. (2005) and Martineau et al. (2018) observed substantial motor-unit enlargement in the STM during disease progression, but these events appeared to be mostly due to nodal sprouts. This is consistent with the absence or disorganization of nerve terminal sprouts (Fig. 10) and the apparent sparing of axonal SCs documented here (Fig. 8).

### Motor-unit type-dependent and independent diversity in PSC properties

Results presented here and previously (Arbour et al., 2015) show (1) that increased mAChR-dependent PSC activity is present long before disease onset, (2) that this alteration is associated with an inability to adopt a phagocytic phenotype upon denervation in presymptomatic animals, and (3) that PSCs’ phenotype is incompatible with NMJ repair in symptomatic animals, regardless of the MU-type they are associated with. However, excitability and repair properties of PSCs associated with vulnerable FF MUs (STM) contrasted with those observed on PSCs associated with less vulnerable FR and S MUs (SOL).

Indeed, PSC Ca^2+^ signaling is affected differentially depending on the MU-type in *SOD1*^*G37R*^ mice, revealing a unique set of complementary changes in PSC activity at FF NMJs in contrast with those reported at FR and S MUs (Arbour et al., 2015). For instance, while maintaining a higher relative mAChR contribution, nerve-induced PSC Ca^2+^ responses on FF MUs were smaller, possibly due to a reduced sensitivity of the purinergic system (Fig. 2 and 4). In contrast, PSC Ca^2+^ responses on FR and S MUs were larger due to an increased activation of mAChRs (FR and S) and an increased sensitivity of the purinergic system (FR, pre-onset stage only) (Arbour et al., 2015). This highlights the importance of the balance regulation of PSCs activity between muscarinic and purinergic receptors. A possible explanation for these opposite changes may be linked to the differential MU-specific changes in NMJ activity occurring in SOD1 mice (Tremblay et al., 2017). Indeed, chronic changes in NMJ activity have been shown to induce plastic changes in PSC properties (Belair et al., 2010).

Furthermore, our results suggest that changes in PSC excitability may only be transient on FF NMJs while they seem to become more pronounced at the pre-onset stage on S and FR MUs (Arbour et al., 2015). These results suggest that the altered PSC excitability may also confer some benefits on NMJ stability (i.e. before it denervates). Alternatively, but not exclusively, transient changes in PSC excitability may reflect longer-lasting changes in PSC function. Further examination of PSC excitability on denervated NMJs in symptomatic SOD1 mice could shed light on this question. Unfortunately, nerve-muscle preparations from symptomatic SOD1 animals are more fragile and PSCs on denervated NMJs seem more vulnerable to the AM loading procedure (Arbour et al., 2015). Future investigation of PSC properties using another approach, such as GCaMP expression (Heredia et al., 2018), may potentially circumvent those limitations.

Moreover, PSC repair properties varied according to the MU type in symptomatic *SOD1*^*G37R*^ animals, where PSC properties on FF NMJs were out of phase with the innervation status while PSCs on FR or S NMJs adopted a less pronounced phenotype. Similarly, the expression of the chemorepellent Semaphorin 3A was selectively increased in FF NMJs following denervation or blockage of neurotransmitter release (De Winter et al., 2006). All these results are consistent with the known vulnerability of FF MUs compared to FR and S MUs in the disease (Frey et al., 2000; Atkin et al., 2005; Pun et al., 2006). Importantly, these differences were not due to STM being further along the progression of the disease than the SOL considering that denervation levels were comparable between the STM and the SOL at this stage. Hence, PSC repair properties reflect MU vulnerability at presymptomatic and symptomatic stages of the disease. Furthermore, the lack of PSC-dependent synaptic repair at FF NMJs could contribute to their increased vulnerability in ALS.

An additional level of complexity arises from the increased heterogeneity in PSC activity (nerve-induced Ca^2+^-responses) and repair properties (Gal-3 expression and process extensions) at individual NMJs in *SOD1* animals. Heterogeneity in PSC responses to synaptic activity could arise from two scenarios. First, a mAChR-independent component of PSC Ca^2+^ responses could be inconsistently decreased amongst PSCs. Second, knowing that PSCs cover exclusive synaptic territories (Brill et al., 2011) and that synaptic activity is altered at the NMJ in ALS (Armstrong and Drapeau, 2013b, a; Rocha et al., 2013; Shahidullah et al., 2013; Arbour et al., 2015; Tremblay et al., 2017), neurotransmitter release could be more heterogeneous between active zones, thus inducing variable levels of Ca^2+^ activity in PSCs depending on the active zones they cover.

Altogether, common pathological alterations in PSC properties are complemented by numerous context-dependent differences that could contribute to, or at least reflect, NMJ dysfunction and MU-vulnerability in ALS.

### Non-cell autonomy at the NMJ and impact of PSC dysfunction on disease progression

Similarly to glial cells in the CNS (Boillee et al., 2006a; Boillee et al., 2006b; Yamanaka et al., 2008; Ilieva et al., 2009; Haidet-Phillips et al., 2011; Kang et al., 2013; Meyer et al., 2014; Ditsworth et al., 2017), results presented here and in recent studies (Carrasco et al., 2016a; Van Dyke et al., 2016), suggest that alterations in PSC properties could contribute to ALS onset and progression. Direct manipulation of mutant SOD1 levels in astrocytes, microglia or oligodendrocyte progenitors directly affected their function and revealed their contribution to the disease. These results raise the question of whether changes in PSCs are directly caused by the expression of the mutant protein or reflect a maladaptive response to the diseased motoneuron. Previous studies evaluating the contribution of mutant SOD1 expression in SCs (both PSCs and axonal SCs) to the disease provided disparate results depending on the dismutase competency of the mutated SOD1 protein (Lobsiger et al., 2009; Turner et al., 2010; Wang et al., 2012).

However, the P0 promoter used in these studies poorly targets PSCs (Lobsiger et al., 2009). Furthermore, we show here that axonal SCs robustly adopted a phagocytic phenotype following denervation in symptomatic *SOD1* mice, while PSCs failed to do so (Fig. 8), suggesting that different types of Schwann cells may be affected differently in the disease. Unfortunately, lack of a known specific PSC promoter has so far prevented a direct manipulation of mutant SOD1 expression in PSCs. Interestingly, Castro et al. (2020) recently suggested a genetic strategy to identify PSCs which could open the possibility to develop new tools to target them selectively. Hence, the contribution of mutant SOD1 expression in PSCs to their dysfunction and to ALS onset and progression remains to be determined.

### Role of Gal-3 in ALS

Numerous studies have evaluated the role of Galectin-3 (MAC-2) in ALS and its usefulness as a potential biomarker for the disease (Zhou et al., 2010). For instance, Gal-3 expression is increased early in the spinal cord of *SOD1*^*G93A*^ mice and ALS patients, in skeletal muscles of *SOD1*^*G86R*^ mice (possibly in axonal SCs based on our results), and rises throughout disease progression (Ferraiuolo et al., 2007; Gonzalez de Aguilar et al., 2008; Zhou et al., 2010; Lerman et al., 2012; Baker et al., 2015). Its expression correlates with microglial activation (Yamanaka et al., 2008; Lerman et al., 2012) and is also inexplicably upregulated in motoneurons during disease progression (Ferraiuolo et al., 2007). These observations have been interpreted as indications that Gal-3 expression could play a detrimental role in disease progression or would at least reflect increasing levels of neuroinflamation.

However, Gal-3 has been implicated in numerous cellular functions which could be protective in ALS (Dumic et al., 2006; Yang et al., 2008) such as phagocytosis in astrocytes (Nguyen et al., 2011; Baker et al., 2015; Morizawa et al., 2017) and myelin phagocytosis in axonal SCs (Reichert et al., 1994). Although still debated, evidence suggest it could play a similar role in microglia (Rotshenker, 2009; Lalancette-Hebert et al., 2012). These results suggest that its upregulation may be linked to increased cellular debris clearance in ALS. Our results suggest that extracellular Gal-3 at the NMJ and in peripheral nerves would exert a beneficial effect on disease progression, being inversely correlated to the accumulation of axonal debris. However, a role of intracellular Gal-3 cannot be cast out as both fractions could not be labeled simultaneously. Lack of debris clearance would delay NMJ reinnervation (Kang and Lichtman, 2013) causing PSCs to gradually abandon parts of the endplate, thus leading to incomplete NMJ reinnervation (Kang et al., 2014). In agreement with this possibility, crossing *SOD1*^*G93A*^ mice to Gal-3 knock-out (*lgal3*^*−/-*^) mice significantly accelerated disease progression (Lerman et al., 2012), further supporting a protective role of Gal-3 in ALS. Overall, these results suggest that the effect of Gal-3 in ALS is context-specific, exerting different adaptive or maladaptive roles in different cell types.

## Conclusion

We show that PSC excitability and repair properties are incompatible with NMJ repair across all MU types, in two *SOD1* ALS mice models, in a manner which reflects their selective vulnerability. These results suggest that compensatory NMJ reinnervation may be defective in ALS due to misguided nerve terminal sprouting and inadequate endplate debris clearance. Further studies aimed at understanding the molecular basis of the diversity and the changes in PSCs in ALS could unravel novel therapeutic targets. Restoring PSCs’ repair capabilities could enhance synaptic reestablishment which could improve motor function in ALS patients.

## Acknowledgements

This work was funded by grants from the Canadian Institutes for Health Research (R.R. MOP-111070), Robert Packard Center for ALS Research (R.R.), Canadian Foundation for Innovation (R.R), and an infrastructure grant from the Fonds Recherche Québec-Santé Leader Opportunity Fund to the GRSNC. É.M. held a doctoral studentship from the ALS Society of Canada and a master’s degree studentship from the Fonds Recherche Québec-Santé. We thank Félix-Antoine Robert for help regarding the statistical analysis.

